# Functional glycoproteomics by integrated network assembly and partitioning

**DOI:** 10.1101/2023.06.13.541482

**Authors:** Matthew E. Griffin, John W. Thompson, Yao Xiao, Michael J. Sweredoski, Rita B. Aksenfeld, Elizabeth H. Jensen, Yelena Koldobskaya, Andrew L. Schacht, Terry D. Kim, Priya Choudhry, Brett Lomenick, Spiros D. Garbis, Annie Moradian, Linda C. Hsieh-Wilson

## Abstract

The post-translational modification (PTM) of proteins by O-linked β-*N*-acetyl-D-glucosamine (O-GlcNAcylation) is widespread across the proteome during the lifespan of all multicellular organisms. However, nearly all functional studies have focused on individual protein modifications, overlooking the multitude of simultaneous O-GlcNAcylation events that work together to coordinate cellular activities. Here, we describe **N**etworking of **I**nteractors and **S**ubstrat**E**s (NISE), a novel, systems-level approach to rapidly and comprehensively monitor O-GlcNAcylation across the proteome. Our method integrates affinity purification-mass spectrometry (AP-MS) and site-specific chemoproteomic technologies with network generation and unsupervised partitioning to connect potential upstream regulators with downstream targets of O-GlcNAcylation. The resulting network provides a data-rich framework that reveals both conserved activities of O-GlcNAcylation such as epigenetic regulation as well as tissue-specific functions like synaptic morphology. Beyond O-GlcNAc, this holistic and unbiased systems-level approach provides a broadly applicable framework to study PTMs and discover their diverse roles in specific cell types and biological states.

## INTRODUCTION

Post-translational modifications (PTMs) allow cells to modulate protein function rapidly and dynamically in response to biological stimuli. O-GlcNAc glycosylation (known as O-GlcNAcylation) is the reversible addition of the monosaccharide β-*N*-acetyl-D-glucosamine to Ser and Thr residues of intracellular proteins. O-GlcNAcylation has been identified on thousands of proteins and functions as a major integrator of cellular signals. Metabolic information from anabolic and catabolic pathways for carbohydrates, amino acids, lipids, nucleotides, and energy storage is combined to produce the precursor, uridine 5’-diphosphate (UDP)-GlcNAc.^1^ Cycling of the modification occurs rapidly in response to many stimuli such as cellular stress,^2^ nutrient availability,^3^ and extracellular signaling.^4^ Accordingly, O-GlcNAcylation regulates numerous essential functions, including transcription,^4^ translation,^5^ metabolism,^6, 7^ protein homeostasis,^8^ and signal transduction.^9, 10^ The broad involvement of O-GlcNAcylation in critical processes, along with its dynamic responsiveness to wide-ranging signals, suggests high-level, global coordination of the modification in cells. Despite this, few studies have examined how multiple O-GlcNAcylation events across the cell are coordinated to orchestrate distinct cellular responses.

Functional studies of O-GlcNAcylation are challenging due to both biological and technical issues. As detection of the labile modification can be difficult, intense efforts have been dedicated to map O-GlcNAcylation sites across the proteome using mass spectrometry.^11^ Despite significant technological advances, methodologically-driven studies often yield datasets that are decoupled from their physiological context, which hinders the selection and prioritization of functionally relevant modification sites. Unlike other PTMs, O-GlcNAc cycling is carried out by only two essential enzymes, O-GlcNAc transferase (OGT) and hydrolase (OGA).^12^ Therefore, traditional methods such as gene silencing or pharmacological inhibition disrupt multiple modifications simultaneously across the proteome. Moreover, no tools exist to modulate O-GlcNAc levels at individual sites on endogenous proteins in cells for gain-of-function studies. As such, functional studies typically require expression of individual glycosylation-deficient proteins via low throughput methods such as gene editing or viral-mediated, stable integration. Given these limitations, nearly all O-GlcNAc sites have been studied in isolation, independently from other sites, and the vast majority of sites remain functionally uncharacterized.

Although single-target studies have been crucial for understanding O-GlcNAc function, such analyses ignore the full breadth of O-GlcNAc activity across the cell and do not address how the composite of multiple modifications works together to direct cellular processes. For example, studies have independently shown that cancer cells alter the O-GlcNAcylation status of c-myc (MYC), phosphofructokinase (PFK1), Tet methylcytosine dioxygenases (TET), and cofilin (CFL1) to reprogram metabolic flux,^13^ gene expression,^14, 15^ and cytoskeletal dynamics,^16^ influencing tumor aggressiveness and metastatic potential presumably in a collective fashion.^15–17^ Moreover, neurons dynamically modulate the O-GlcNAcylation status of proteins from the synapse to the nucleus in response to excitatory and inhibitory stimuli.^4, 7, 18, 19^ In these cases, O-GlcNAc cycling occurs on multiple time scales to transiently regulate protein activity, localization, and stability, while also influencing long-term genetic reprogramming through transcription factor recruitment and epigenetic markers.^4, 20, 21^ Finally, environmental cues such as nutrient availability and insulin signaling can initiate distinct O-GlcNAcylation responses across different tissues such as liver and adipose tissue.^9, 22, 23^ Thus, new methods are needed to parse out the diverse effects of O-GlcNAcylation in a given biological context, while providing a holistic understanding of how key glycosylation events work together to drive specific cellular outcomes.

The ‘adaptor protein hypothesis’ has been proposed to explain the diverse cellular activities of OGT.^1^ This hypothesis posits that adaptor proteins may serve as spatial and temporal regulators of O-GlcNAcylation by targeting OGT to cellular substrates, similar to E3 ligases that target substrates for ubiquitination and subsequent degradation.^24^ We envisioned that OGT interacting proteins might provide an organizational framework for understanding the activity, specificity, and regulation of O-GlcNAcylation within cells. Thus, the unique set of OGT interactors could reveal the concurrent functions of the modification in a particular biological setting. However, such an analysis would require new robust, quantitative methods for profiling both OGT interactors and substrates, as well as powerful computational methods for integrating and functionally parsing the resulting large datasets.

Here, we describe the **N**etworking of **I**nteractors and **S**ubstrat**E**s (NISE) approach for comprehensive, systems-level analyses of PTMs. As applied to O-GlcNAc, the NISE method first determines important molecular components of O-GlcNAc activity by identifying OGT interacting proteins (**Figure 1A**) and O-GlcNAcylated substrates (**Figure 1B**) for a given cellular context. These elements are then combined using global protein-protein interaction (PPI) datasets to produce a functional O-GlcNAcylation network that is partitioned into process-specific subnetworks by unsupervised clustering (**Figure 1C**). Notably, this integrated approach provides new insights that cannot be obtained from the individual interactome or O-GlcNAcome datasets alone. Together, the NISE approach offers a holistic, proteome-wide view of OGT activity for a given biological state, which serves as a powerful framework for generating functional hypotheses and identifying targets for direct interrogation.

**Figure 1.**
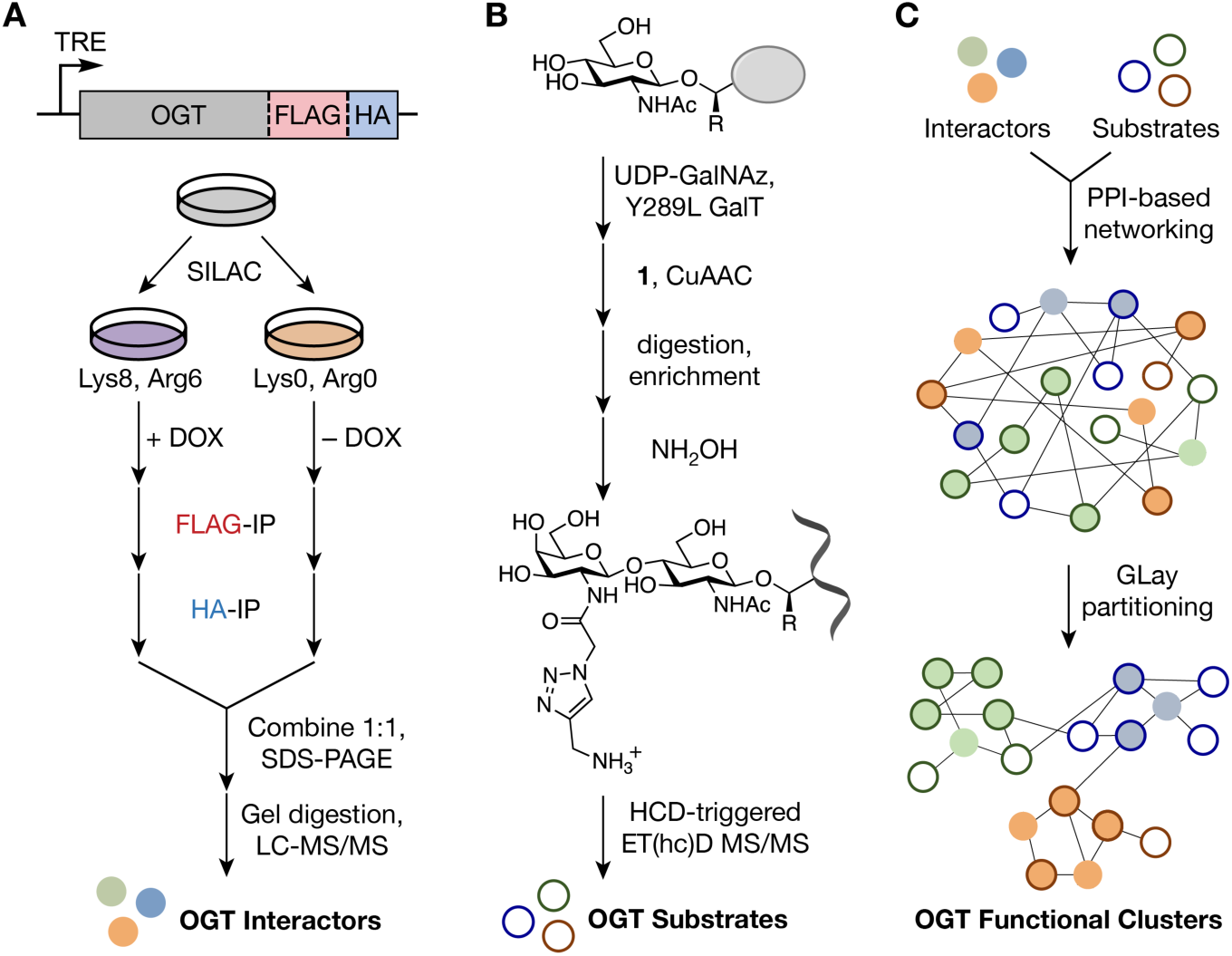
Networking of Interactors and Substrates (NISE). (A) OGT interactors are isolated by tandem affinity purification (TAP) of doxycycline (DOX) inducible, dual-tagged OGT using stable isotope labeling by amino acids in cell culture (SILAC) for quantification and identified by mass spectrometry. (B) OGT substrates are biotinylated by selective chemoenzymatic labeling of the O-GlcNAc moiety and copper-catalyzed azide-alkyne cycloaddition (CuAAC) with a chemically cleavable biotin tag (**1**, see Fig. 2E), followed by tryptic digestion, affinity purification of the labeled peptides, and mass spectrometry-based site identification. R = H for serine or OH for threonine. (C) Networks using context-specific OGT interactor and substrate datasets are constructed using established protein-protein interactions, followed by community-based partitioning to assemble functionally related subclusters.

We first demonstrated the approach by identifying 519 robust OGT-protein interactions and 1,764 O-GlcNAc modification sites in HEK293T cells and producing a network that revealed both known and novel roles for O-GlcNAcylation in protein translation, epigenetic regulation, mRNA splicing, and other cellular processes. By incorporating additional bioinformatics data regarding protein domains, amino acid motifs, and other PTMs into this network, we created an information-rich resource that gives new insights into protein and pathway regulation by O-GlcNAcylation. Exploiting the unique connectivity provided by NISE, we also identified putative OGT adaptor proteins and provide direct evidence for both positive and negative regulation of site-specific O-GlcNAcylation by a single adaptor protein. Additionally, we expanded our approach in vivo (iNISE) through the generation of a novel mouse model expressing endogenously dual tagged OGT. Using this model, we identified 993 and 643 high-confidence OGT interactors in healthy brain and liver tissue, respectively, as well as a total of 2,785 non-redundant O-GlcNAc sites, which were integrated to produce tissue-specific iNISE networks. Network comparisons revealed conserved, core activities of O-GlcNAc across cells and tissues, including histone modification, mRNA processing, and actin dynamics, as well as unique, tissue-specific functions such as metabolism and synaptic structure. Together, these studies provide a global framework and comprehensive databases for uncovering OGT specificity and O-GlcNAc function with unprecedented depth and scale.

## RESULTS

### Identification of statistically validated, native OGT-interacting proteins

To build O-GlcNAc networks for NISE analysis, we first required comprehensive, robust, and general methods to identify context-specific OGT-interacting proteins. Although OGT interactors have been reported using both targeted and global PPI approaches,^25–28^ previous efforts have relied on yeast two-hybrid assays^29^ and protein microarrays,^27^ which were performed under non-native conditions that do not account for cell state- or cell type-specific interactions. Affinity pulldowns from living cells better contextualize OGT function in physiologically relevant settings and can capture both direct and indirect interactors, revealing larger protein complexes that encompass OGT.^26, 28, 30^ However, co-immunoprecipitation methods have generally been performed in unicate, with arbitrary affinity cutoffs and limited information on statistical reproducibility. Thus, we combined tandem affinity purification-mass spectrometry (TAP-MS), a well-established method for high-throughput, native-state interactomics,^26, 31, 32^ with stable isotope labeling by amino acids in cell culture (SILAC) and spectral counting for accurate protein quantification, to generate the first statistically robust, native OGT interactome.^33, 34^

Full-length OGT with dual FLAG and HA epitope tags (OGT-FH; **Figure 1A**) was stably expressed in HEK293T cells under the tetracycline-responsive element (TRE) promoter using lentiviral vectors. This system allowed for uniform, controllable levels of OGT-FH (**Figures S1A and S1B**) via the addition of doxycycline (DOX). OGT-FH and its associated proteins were then subjected to two sequential rounds of immunoprecipitation (**Figures 1A and 2A**). Several known OGT interactors were pulled down and detected by western blotting, including retinoblastoma binding protein 5 (RBBP5),^35^ nuclear pore glycoprotein p62 (NUP62),^36^ WD repeat-containing protein 5 (WDR5),^35^ and host cell factor 1 (HCFC1)^35^ (**Figure 2B**), validating the TAP procedure. For quantitative interactome analyses, we compared proteins immunoprecipitated from DOX treated, OGT-FH-expressing cells with those immunoprecipitated from vehicle-treated, non-OGT-FH-expressing cells using both SILAC ratios and spectral counting for all proteins. This approach was performed in triplicate, which allowed us to limit potential false positives and provide a statistically validated set of OGT-interacting proteins. We identified 519 high-confidence OGT interactors with a corrected *P*-value of < 0.05 from either quantitation method (**Figure 2C**; **File S1**). Within the dataset, only 64 (12%) were previously described OGT interactors annotated within the PPI databases BioGRID^37^ and IntAct.^38^ All interactors were found in two or more independent replicates, and the vast majority (498, 96%) were observed in all three replicates (**Figure 2D**), highlighting the robustness of our OGT interactome data and TAP-MS platform. Importantly, many high-confidence interactors have been associated with major advances in the O-GlcNAc field, such as host cell factor C1 (HCFC1),^26, 39^ BRCA1 associated protein 1 (BAP1),^40^ and ten-eleven-translocation 2 (TET2).^41^ Moreover, we identified 455 new interactors, underscoring the importance of conducting targeted, comprehensive OGT interactome analyses. Among the new interactors were DNA damage-binding protein 1 (DDB1), mitotic checkpoint protein BUB3 (BUB3), and serine/threonine-protein kinase WNK1 (WNK1), which play critical roles in DNA damage repair,^42^ mitotic spindle assembly,^43^ and ion homeostasis,^44^ respectively.

**Figure 2.**
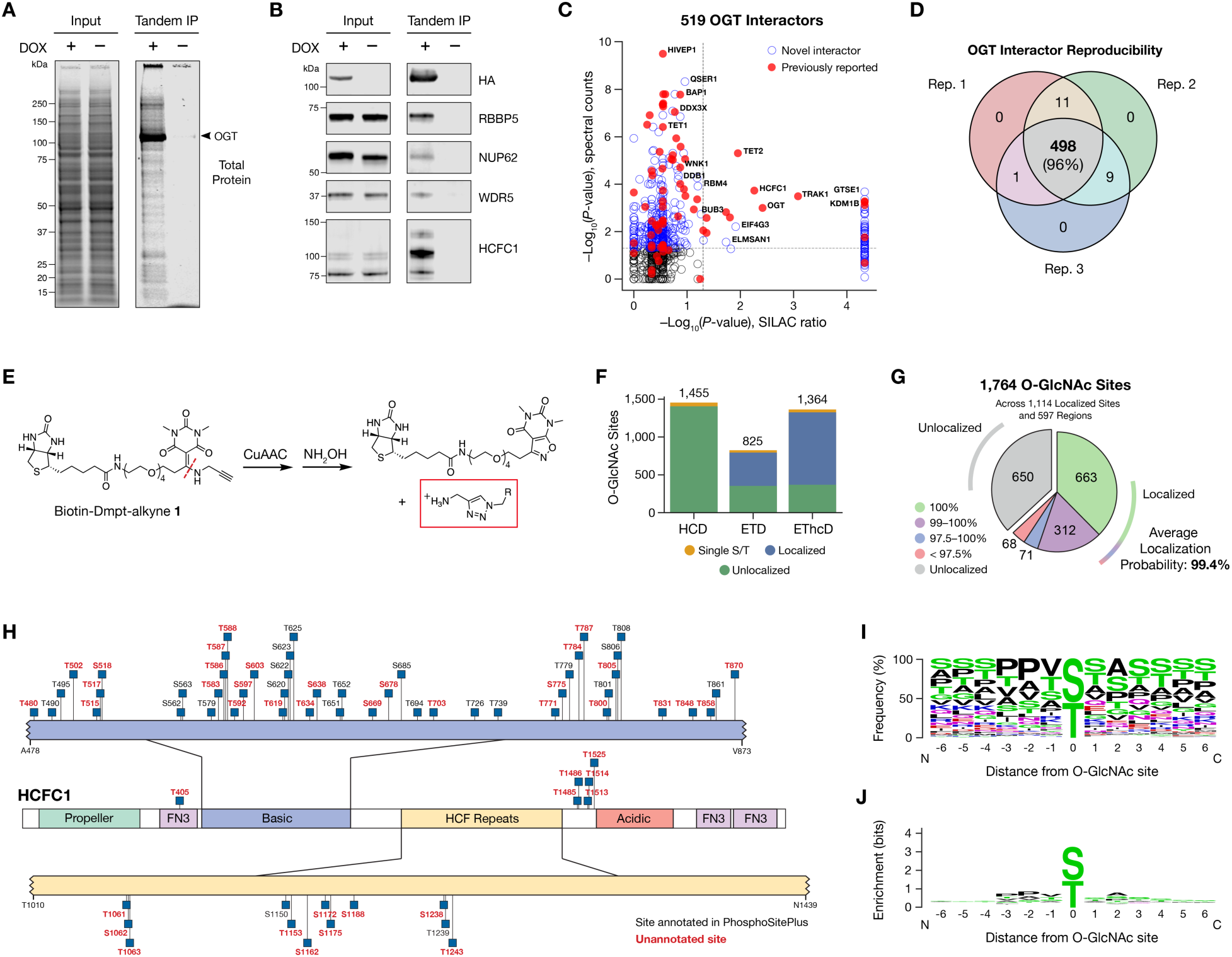
Generation of context-specific OGT interactor and substrate datasets. (A) Silver-stained gel profiles of cell lysates and purified OGT interactors after TAP using HEK293T cells stably expressing OGT-FLAG-HA. (B) Validation of the TAP method via western blotting for known OGT interactors. (C) Scatterplot of statistically validated OGT interactors identified from HEK293T cells. (D) Venn diagram of putative OGT interactors identified across three independent replicates. (E) Chemical structure and hydroxylamine-mediated release of cleavable biotin tag **1**. Dmpt = *N*,*N*’-dimethylpyrimidinetrione, R = labeled O-GlcNAcylated peptide. (F) Quantification of localized and unlocalized O-GlcNAc sites identified by MS^2^ fragmentation methods. For localized sites, the exact site of the modification was determined with a calculated probability of confidence in the assignment (see Fig. S2). (G) Distribution of localization probabilities of O-GlcNAc sites identified from HEK293T cells. (H) Annotation of localized O-GlcNAc sites identified on host cell factor 1 (HCFC1), with sites not annotated in PhosphoSitePlus shown in red. (I) Frequency plot of amino acid residues surrounding localized O-GlcNAc sites. (J) Enrichment plot of amino acid residues surrounding localized O-GlcNAc sites.

### Proteome-wide mapping of protein O-GlcNAcylation sites

To generate a comprehensive database of OGT substrates in HEK293T cells, we expanded our chemoenzymatic labeling (CL) approach to identify sites of O-GlcNAcylation across the proteome.^45, 46^ This method employs a mutant galactosyltransferase (Y289L GalT) to append the non-natural monosaccharide β-*N*-azidoacetyl-D-galactosamine (GalNAz) onto O-GlcNAc modified proteins as a handle for downstream functionalization and enrichment. CL is advantageous over other methods such as metabolic labeling^47^ due to its high specificity toward O-GlcNAcylated substrates, quantitative labeling efficiency, and lack of off-target, nonenzymatic cysteine labeling.^46, 48^ Furthermore, the CL method can be applied to any biological materials because it does not require the treatment of live samples with exogenous, non-natural sugars, which can also alter physiological glycosylation levels. As such, CL enables individual O-GlcNAc sites to be readily identified and quantified across different cells and tissues.

To enhance the capabilities of our CL approach further,^45, 49^ we developed a new chemically cleavable linker for improved capture-and-release of modified targets. Biotin-Dmpt alkyne **1** (**Figures 2E and S1C**) incorporates an *N,N’-*dimethylpyrimidinetrione (Dmpt) group that exhibits higher chemical and thermal stability than the 1-(4,4-dimethyl-2,6-dioxocyclohex-1-ylidene)ethyl (Dde) scaffold used in our previous generation linker (**Figure S1D**).^45^ The Dmpt group also provides a versatile tag for targeted MS enrichment workflows, as it is stable during proteolytic digestion with trypsin and is removed quantitatively with dilute solutions of alpha nucleophiles such as hydroxylamine. Upon cleavage, the resulting primary amine aids in glycopeptide fragmentation via electron transfer dissociation (ETD) by increasing the peptide charge density.^50^ Moreover, the tagged peptides produce a signature ion of 300.1 *m/z* during higher-energy collisional dissociation (HCD), which allows for the rapid, definitive identification of O-GlcNAcylated peptides.^45, 51^

We developed an efficient MS method that capitalized on this unique signature ion and targeted the tagged glycopeptides for sequencing and O-GlcNAc site identification. Detection of the 300.1 *m/z* reporter ion by HCD was used to trigger fragmentation of the precursor ion by ETD or ETD with HCD supplemental activation (EThcD), leading to faster instrument cycle times and avoiding the time-intensive fragmentation of unmodified peptides. Two biological replicates were processed in parallel, and each replicate was split into two separate LC-MS/MS runs using HCD triggered ETD or HCD-triggered EThcD to maximize the number of site identifications. Overall, HCD identified the most O-GlcNAcylated peptides, followed by EThcD and then ETD (**Figure 2F**). However, HCD also led to neutral loss of the glycan residue for nearly all peptides as expected.^46, 52^ Notably, we observed that ETD and EThcD each identified unique, non-overlapping sets of sites (**Figure S1E**), underscoring the importance of a combined fragmentation approach to maximize site identification and localization.

To map and count the O-GlcNAc sites in our dataset, we developed a novel computational program that localized the modification to exact sites or short peptide regions (**Figure S2**). Unlike standard methods for counting PTM sites, this approach also eliminated duplicate sites from missed cleavages or different peptide charge states to provide an accurate total number of non redundant O-GlcNAc sites, rather than the commonly reported total number of nonidentical modified peptides. This general strategy is directly applicable to any variable peptide modification, including PTMs such as phosphorylation, acetylation, and ubiquitination. Together, the analysis of HEK293T cells yielded 1,764 unique O-GlcNAcylated sites (**File S2)** across 646 proteins (**File S3**), with a false discovery rate of 1%. Notably, our improved MS ionization methods produced highly efficient peptide fragmentation without loss of the glycan moiety, allowing us to localize the O-GlcNAc modification to a single Ser/Thr residue with >99% probability for the majority of sites (**Figure 2G**). For example, we were able to definitively assign 66 of the 72 total O-GlcNAc sites and regions that we detected on HCFC1, including 47 sites that were not previously annotated in the PhosphoSitePlus database (**Figure 2H**).^53^ Frequency (**Figure 2I**) and enrichment analysis (**Figure 2J**) of all localized sites using WebLogo^54^ revealed the lack of a consensus sequence for OGT as expected. We found a slight preference for small hydrophobic residues near the modification site, including proline at the -3 and -2 positions, as reported previously,^55, 56^ as well as higher frequencies of both proximal and distal Ser and Thr residues.

### Generation and validation of functionally partitioned O-GlcNAc networks

We next sought to integrate the large datasets of OGT interactors and substrates from HEK293T cells to gain insights into how multiple O-GlcNAcylation events may be coordinated to regulate cellular function. First, we connected our datasets of interactors and substrates (or “nodes”) through defined, physical interactions. Known PPIs between OGT interactor–interactor pairs or OGT interactor–substrate pairs were identified from the BioGRID or IntAct databases and used as undirected “edges” between nodes, resulting in a functional O-GlcNAc network that was visualized using Cytoscape.^57^ To partition the network into individual clusters for further analysis, we then employed the community clustering algorithm GLay.^58–60^ This program scores edge betweenness for node groups to extract local communities from a larger network and has been previously used to visualize and partition interaction networks into functionally relevant clusters.^59, 61–63^ Importantly, the GLay algorithm is inherently agnostic to protein identity, which provides a functionally unbiased approach to network partitioning. Partitioning of the complete network revealed ten, highly intra-connected communities, five of which had more than ten proteins (**Figure 3A**). To assess whether the communities contained functionally related nodes, we performed a Gene Ontology (GO)^64^ analysis using biological process terms. Grouping of the significant GO terms using ClueGO^65^ revealed distinct cellular functions within the five major subnetworks: 1) posttranscriptional regulation; 2) chromatin organization and remodeling; 3) nuclear and organelle transport and cytoskeleton organization; 4) RNA/DNA synthesis and catabolism; and 5) mRNA splicing and translational initiation (see **Figure S3A** for full annotation of all clusters).

**Figure 3.**
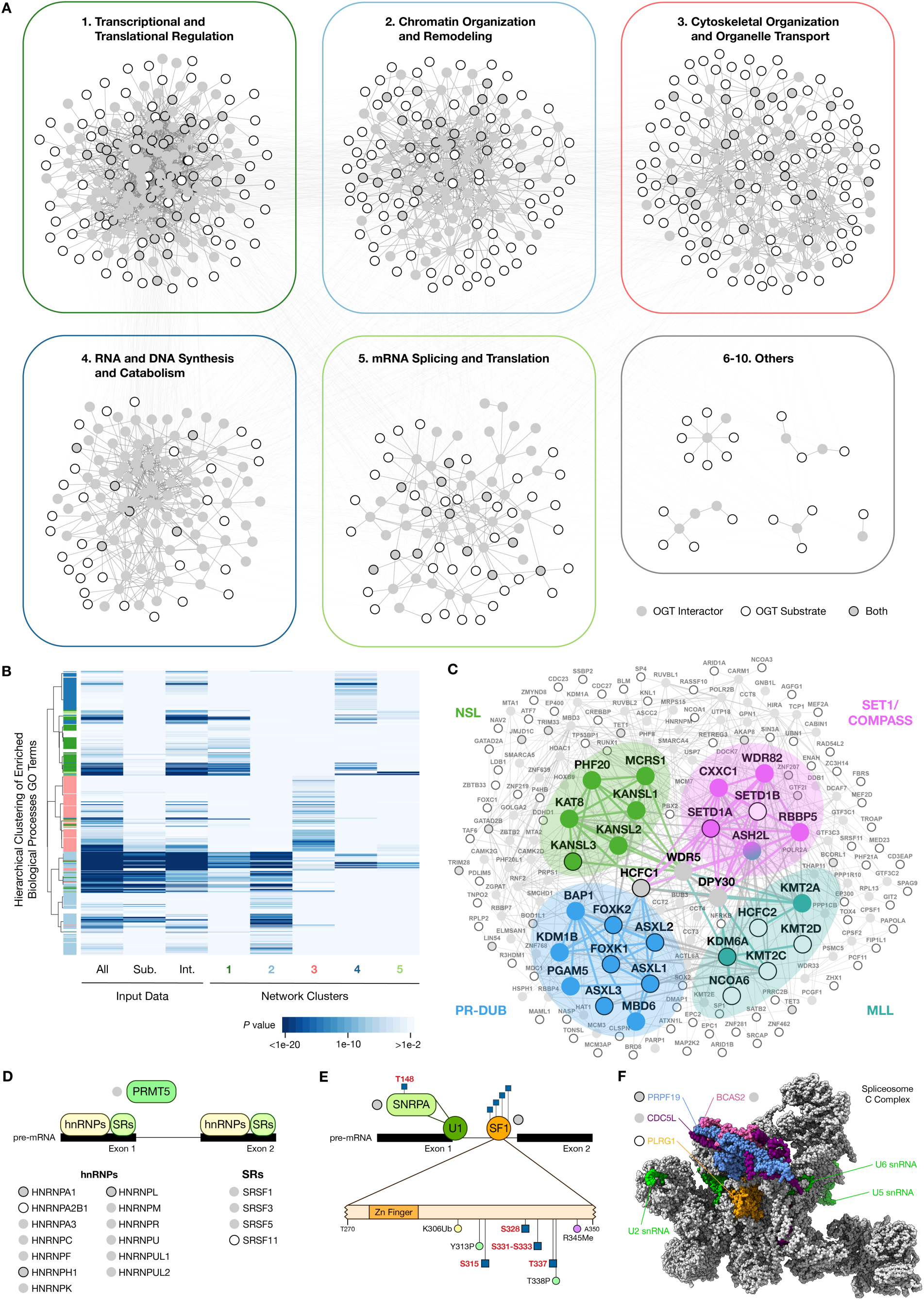
NISE reveals diverse potential regulatory roles for the O-GlcNAc modification. (A) O-GlcNAc functional network from HEK293T cells with communities annotated by enriched biological process GO terms. (B) Hierarchical clustering of enriched biological process (BP) GO terms from initial interactor and substrate datasets and individual GLay communities demonstrates efficient partitioning of biological functions. (C) Annotation of subcluster 2 highlighting four protein complexes (NSL, SET1/COMPASS, PR-DUB, and MLL) involved in epigenetic regulation of gene expression. (D) Schematic of identified OGT interactors and substrates involved in intron retention and pre mRNA stabilization. (E) Schematic of identified OGT interactors and substrates within the spliceosome early (E) complex, including SNRPA within the U1 small nuclear ribonucleoprotein (snRNP) and SF1. Ub = ubiquitination, P = phosphorylation, Me = methylation. (F) Schematic of identified OGT interactors and substrates within the Prp19/Nineteen complex (NTC) of the catalytic (C) spliceosome complex. Structure from PDB 5YZG.

To evaluate the effectiveness of the GLay partitioning, we compared the biological process GO terms that were significantly enriched (*P* < 0.01) in the interactor or substrate datasets alone, the combined dataset, and within each partitioned community. Hierarchical clustering based on co-enrichment demonstrated that the GO terms were partitioned well into individual clusters, with relatively few terms distributed across multiple clusters (**Figure 3B**). Moreover, nearly all GO terms from the original datasets were found to be enriched within the final groupings, suggesting no significant loss of functional data. Similar results were obtained through analysis of molecular function (**Figure S3B**) and cellular component (**Figure S3C**) GO terms, Kyoto Encyclopedia of Genes and Genomes (KEGG) pathways (**Figure S3D**),^66^ Reactome pathways (**Figure S3E**),^67^ WikiPathways (**Figure S3F**),^68^ and Comprehensive Resource of Mammalian Protein Complexes (CORUM) groups (**Figure S3G**),^69^ highlighting the power of the NISE approach to reveal the central functions of O-GlcNAc and delineate their molecular components into individual, biologically-related communities (see also **File S4**).

Importantly, NISE uncovered functional annotations within the individual clusters that were absent from analyses of the interactor, substrate, or combined datasets before GLay partitioning (**Figure S3H**, **File S5**). For example, Cluster 1 was enriched for previously unidentified pathways involved in Toll-like receptor (TLR), tumor necrosis factor (TNF), interleukin-17 (IL17), and mitogen-activated protein kinase (MAPK) signaling, as well as GO terms associated with Fragile X syndrome. In Cluster 2, we identified new potential O-GlcNAc pathways related to HOX gene activation, pre-mRNA processing, and Cushing syndrome. In-depth interrogation of GO terms within Cluster 3 revealed significant enrichment of intracellular transport and cytoskeletal organization pathways. Although these are well-studied activities of O-GlcNAc,^70–74^ they were not observed in the individual or combined datasets before partitioning. Cluster 3 also indicated novel functional links between O-GlcNAc and membrane trafficking, Hippo signaling, and sterol regulator element-binding protein (SREBP) signaling pathways, whereas Cluster 4 showed enrichment of polymerase switching, DNA synthesis, and mismatch and nucleotide excision repair pathways. Finally, Cluster 5 showed enrichment for components of the spliceosome and transcription export complex. In total, our NISE-based analysis uncovered 145 biological process, 20 molecular function, and 40 cellular component GO terms, as well as 9 KEGG, 66 Reactome, 9 WikiPathways, and 43 CORUM complexes that emerged only after network clustering (**File S5**).^66–69^

The functional annotations revealed by NISE provide critical insights into the primary functions of O-GlcNAc for a given cell type or state. For example, Cluster 2 (chromatin organization and remodeling) was enriched for 44 of the 82 CORUM complexes in the network (**Figure S3G; File S4**), including the histone H3K4 methyltransferase complexes SET1/COMPASS^75, 76^ and mixed lineage leukemia (MLL),^77, 78^ the polycomb repressive deubiquitinase (PR-DUB) complex,^35^ and the nonspecific lethal (NSL) histone acetyltransferase complex^79, 80^ (**Figure 3C**). These complexes suggest functional roles for O-GlcNAcylation in the regulation of three major epigenetic histone marks. Consistent with these findings, recent studies have demonstrated that OGT and O-GlcNAcylation contribute to SET1-mediated histone methylation,^75^ MLL stability,^77^ and NSL-mediated histone acetylation.^79^ O-GlcNAcylation was also shown to coordinate transcriptional repression by the PR-DUB complex in *Drosophila*,^81^ and our NISE network indicates that this activity is likely an important, conserved function in HEK293T and other mammalian cells. Moreover, the wealth of site-specific and PPI information in other proteins surrounding the canonical members of these complexes within our network suggests that O-GlcNAc may play broader roles in coordinating the overall activities of these histone-modifying complexes.

In addition, Cluster 5 revealed previously uncharacterized roles for O-GlcNAcylation in RNA splicing. Consistent with this novel function, pharmacological inhibition of OGT has been reported to globally alter detained intron splicing.^82^ Notably, our network contains several splicing factors shown to be differentially phosphorylated upon OGT inhibition, thereby providing possible molecular mechanisms for this intron splicing activity. For example, we detected OGT interactions with known intron retention mediators, including protein arginine N-methyltransferase 5 (PRMT5), numerous heterogeneous ribonucleoprotein particles (hnRNPs), and serine/arginine-rich splicing factors SRSF1, SRSF3, and SRSF5 (**Figure 3D**). Moreover, we discovered O-GlcNAcylation sites on the phosphoprotein SRSF11, which may indicate direct regulation of this factor by proximal O-GlcNAcylation and phosphorylation. Beyond its role in intron splicing, we found evidence that O-GlcNAcylation may alter core and auxiliary spliceosome components to regulate the splicing machinery more broadly (**File S6**).^83^ For example, we found that the E complex components U1 small nuclear polypeptide A (SNRPA) and splicing factor 1 (SF1), which are involved in the initial recognition of splicing sites and commit pre-mRNA to the splicing pathway, act as both OGT interactors and substrates (**Figure 3E**). Interestingly, multiple O-GlcNAcylation and other PTM sites occur on SF1 proximal to its zinc finger domain, suggesting an intricate mechanism of PTM based regulation that may alter SF1 binding to branch point sequences within introns. We also found that OGT interacted with the U2 splicing factor B3 subunits SF3B1, SF3B2, SF3B3, and SF3B4, indicating a potential role for OGT in the formation of the spliceosome A complex. Finally, our network revealed that multiple OGT interactors and substrates comprise the Prp19/Nineteen complex (NTC), a core component of the catalytic spliceosome C complex that is required for intron removal (**Figure 3F**). Together, these data exemplify how our holistic network-based approach can be used to provide deeper insights into established functions as well as discover new activities for O-GlcNAcylation.

### NISE Analysis of PTM Crosstalk and PPI Modulation

O-GlcNAcylation can provide exquisite control over protein activity by influencing other PTMs on the same or nearby amino acid residues.^4, 84–86^ For example, hyperglycemia-induced O-GlcNAcylation of CREB-regulated transcription coactivator 2 (CRTC2) at S70 and S171 directly blocks inhibitory phosphorylation by AMP-activated protein kinase (AMPK) to induce hepatic gluconeogenesis.^87^ To identify new instances of crosstalk between O-GlcNAc and other PTMs, we incorporated PTM databases into our network structure. Integration of PTM information from the PhosphoSitePlus database showed a striking number of proximal PTMs (**Figure 4A; File S7**). Roughly 25% of the 1,101 localized O-GlcNAc sites that we identified within canonical SwissProt protein sequences directly overlapped with annotated phosphorylation sites, and nearly 65% of the sites resided within 10 amino acid residues of a phosphorylation site. For example, dataset integration revealed the well-documented regulatory O-GlcNAcylation site within the transactivation domain of signal transduction and activator of transcription 3 (STAT3) at T717 (**Figure 4B**). This site has been demonstrated to negatively modulate phosphorylation at neighboring Y705 and S727 residues, which are critical for STAT3 transcriptional activity during inflammation and neural stem cell differentiation.^88, 89^ Additionally, we detected O-GlcNAcylation directly at the S727 phosphorylation site, which may provide further, unexplored mechanisms for regulating STAT3 function. Beyond phosphorylation, hundreds of O-GlcNAc sites were also found within 10 amino residues of acetylation, methylation, or ubiquitination sites, indicating that PTM crosstalk with O-GlcNAc may be an underappreciated, general feature of cell signaling and protein regulation.

**Figure 4.**
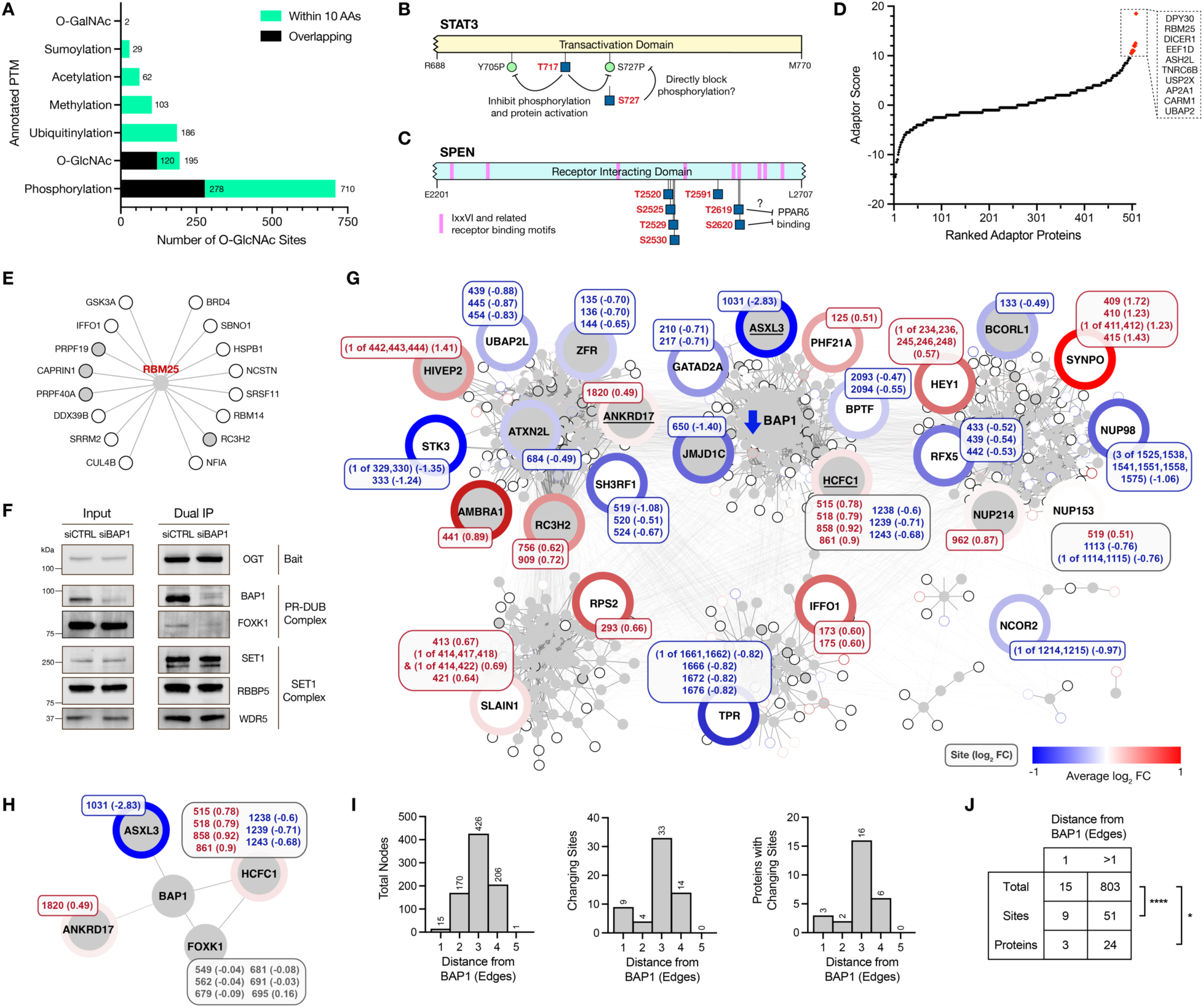
Integration of existing datasets into NISE-generated networks uncovers potential molecular mechanisms of O-GlcNAc regulation. (A) Quantification of post-translational modification (PTM) sites within the PhosphoSitePlus database that overlap with (black) or are proximal to (green) O-GlcNAc sites within the functional network. (B) Schematic of potential STAT3 regulation by PTM crosstalk between known phosphorylation sites (P) and newly identified O-GlcNAc sites. (C) Schematic of potential SPEN regulation by PPI disruption via O-GlcNAcylation of receptor binding motifs. (D) Ranking of potential OGT adaptor proteins within the network. The top ten adaptor proteins are shown. (E) Extracted network of potential OGT adaptor RBM25 and connected OGT substrates. (F) Validation of selective OGT decoupling from the PR-DUB complex via western blotting of PR-DUB and SET1 complex components. (G-H) Site-specific quantification of network-wide and BAP1-adjacent O-GlcNAc site occupancies upon knockout of the OGT interactor BAP1. Only sites with significantly changing occupancies are shown. Log2-fold changes for individual sites are reported as insets on each node, and the node outline color represents the average log2-fold change of all sites on the protein. (I-J) Quantification of edge distances from BAP1 for all nodes, individual changing sites, and proteins with changing sites. * *P* < 0.05, **** *P* < 0.0001; Fisher’s exact test.

Another major mechanism by which O-GlcNAcylation alters protein function is by modifying critical allosteric or protein-protein interfaces to promote or inhibit protein-protein interactions.^13, 90^ Because proteome-wide information regarding the exact amino acid residues found at PPIs is scarce and poorly annotated, we integrated protein domain, motif, region, and amino acid repeat data from the UniProt database^91^ into the network (**File S8**) and examined whether the identified O-GlcNAc sites occurred within protein regions implicated in PPIs. This approach yielded numerous O-GlcNAc sites that overlapped with canonical binding motifs and protein-specific regions identified as protein-protein interfaces. For example, we found seven localized O-GlcNAcylation sites within the nuclear receptor interaction domain (RID) of Msx2 interacting protein SHARP (SPEN), including two sites, T2619 and S2620, that directly overlap with a conserved IxxVI binding motif (**Figure 4C**).^92^ Thus, O-GlcNAcylation of SPEN may alter its association with nuclear receptors such as the metabolic regulator peroxisome proliferator activated receptor delta (PPARδ).^93^ Our analysis also uncovered O-GlcNAcylation of the autophagy regulator serine/threonine-protein kinase ULK1 (ULK1) at S409 in a region that mediates its interaction with the GABA type A receptor-associated protein (GABARAP), which promotes ULK1 kinase activity and autophagosome formation upon starvation.^94^ Finally, we identified a single O-GlcNAc site on homeodomain-interacting protein kinase 1 (HIPK1) at T1001 within a region required for tumor suppressor p53 (TP53) binding that has been suggested to enhance TP53 activation and limit tumor cell growth.^95, 96^ These examples underscore the power of the NISE approach to reveal new, readily testable hypotheses regarding mechanisms of O-GlcNAc function.

### NISE Analysis to Identify Adaptor Proteins for Selective O-GlcNAc Modulation

The context-dependent mechanisms by which OGT activity is regulated toward specific cellular substrates remain poorly understood. Adaptor proteins have been proposed to tether OGT in proximity to particular substrates and selectively modulate their O-GlcNAcylation status in response to diverse signals. Because the proteome-wide networks generated by NISE are built upon physical interactions between proteins, we hypothesized that NISE could be used to identify putative adaptor proteins and investigate the adaptor protein hypothesis in an unbiased, proteome-wide manner. Toward this end, we developed a program to rank OGT interactors based on their relative connectivity. Interactors were scored positively (+1) for each connection or edge to an OGT substrate and penalized (−0.5) for each connection to another OGT interactor. This approach facilitated the identification of OGT interactors that were highly connected to multiple substrates, without over-selecting for members of large protein complexes (**Figure 4D; File S9**). Among the top hits were protein dpy-30 homolog (DPY30) and Set1/Ash2 histone methyltransferase complex subunit ASH2 (ASH2L), which are core scaffolding components of the SET1/COMPASS and mixed lineage leukemia (MLL) complexes (**File S9**).^97–99^ Given the established roles of O-GlcNAc in transcriptional regulation, recruitment of OGT to active promoter and enhancer regions by DPY30 and ASH2L may control associated epigenetic machinery in genomic regions of active transcription. Furthermore, we identified the histone-arginine methyltransferase CARM1 (CARM1) within the top 10 putative OGT adaptors. CARM1 also plays an important role in transcriptional and epigenetic regulation and was previously proposed to function as an OGT adaptor protein.^29^ Beyond transcriptional control, other highly-ranked putative adaptors suggested multiple roles for targeted O-GlcNAcylation in the post-transcriptional regulation of protein expression. These hits included the RNA exon junction complex component, RNA-binding protein 25 (RBM25), which engages spliceosome components and OGT substrates PRPF19, PRPF40A, and SRRM2 (**Figure 4E**), and the RNA-induced silencing complex component, endoribonuclease Dicer (DICER1).

The existence of multiple OGT adaptor proteins could enable O-GlcNAcylation of specific substrates to be decoupled from one another and modulated without affecting global O-GlcNAc levels. Despite strong interest in modulating specific, therapeutically-important OGT substrates such as tau and ɑ-synuclein, no methods exist to selectively alter the O-GlcNAcylation of endogenous protein targets. To test whether the NISE approach could guide the identification of potential adaptor proteins and predict the responsiveness of O-GlcNAc modification sites across the proteome, we examined the protein deubiquitinase BAP1, a component of the PR-DUB complex (**Figure 3C**). BAP1 was connected to several OGT substrates in the network, including forkhead box protein K1 (FOXK1) and additional sex combs-like protein 3 (ASXL3), suggesting it may function as an adaptor protein. Consistent with the notion of decoupled OGT-adaptor protein-substrate complexes, siRNA knockdown of BAP1 in HEK293T cells disrupted the association of OGT with FOXK1, but not with components of the SET1/COMPASS complex, such as SET1, RBBP5, and WDR5 (**Figure 4F**). These results indicate that the OGT-BAP1 complex may function independently from the OGT-SET1/COMPASS complex.

We next investigated whether OGT activity toward substrates proximal to BAP1 in the network was decoupled from other substrates. For this, we knocked out BAP1 in HEK293T cells using CRISPR/Cas9 (**Figure S4A**) and evaluated O-GlcNAc levels at sites across the proteome in wild-type and BAP1-knockout HEK293T cells. To quantify relative O-GlcNAc site occupancy, an MS workflow using tandem mass tag (TMT)-synchronous precursor selection (SPS)-MS^3^ was developed (**Figure S4B**). Because TMT labeling also modifies the primary amine installed on the O-GlcNAc residue after purification with our biotin-Dmpt-alkyne linker, we took advantage of three new diagnostic ions at 732.4, 547.3, and 529.3 *m/z* produced upon HCD fragmentation (**Figure S4C**) to trigger both ETD and EThcD fragmentation, as well as SPS-MS^3^ quantification, further reducing overall instrument cycle times. A total of 1,087 sites across 365 unique proteins was identified and quantified from top speed (5 sec) and top 10 data-dependent runs (**Figure S4D**) over three biological replicates (**File S10**). Here, where every precursor was fragmented by both ETD and EthcD in the same run, we found that ETD and EthcD identified similar numbers of sites (**Figure S4E**) and that these sites were relatively distinct (**Figure S4F**). We also determined and tracked the relative occupancy of 935 (86.0%) of these sites across 290 proteins after MS quantification and normalization for protein expression levels. The fold-change values at each site were averaged across the top speed and top 10 runs, and significant differences were determined by a limma-moderated *t* test (*P* < 0.05).

Notably, only a small fraction of the quantified sites (6.8%; 60 sites across 27 proteins) in the network underwent significant changes upon BAP1 knockout (**Figures 4G and 4H**), and these changes were not randomly distributed throughout the network. Instead, we found that the frequency of changing sites and proteins was significantly enriched at nodes directly connected to BAP1 (**Figures 4I and 4J**; sites: *P* < 0.0001, proteins: *P* = 0.0172, Fisher’s exact test). For example, we observed an 85.9% decrease in O-GlcNAcylation at S1031 of ASXL3, although total ASXL3 protein levels were not quantifiable by MS or western blotting analysis. Interestingly, we also found an increase in O-GlcNAcylation of the BAP1 interactor ankyrin repeat domain containing protein 17 (ANKRD17) at S1820 upon BAP1 knockout, and both increases (4 sites) and decreases (3 sites) at distinct sites on the BAP1 interactor HCFC1, suggesting that adaptor proteins such as BAP1 can both activate and inhibit site-specific O-GlcNAcylation of certain substrates. Changes in site occupancy on HCFC1 also covaried in specific regions, with increases clustered at neighboring residues T515/S518 and T858/T861 and decreases at S1238/T1239/T1243. Conversely, O-GlcNAc site occupancy on the direct BAP1 interactor FOXK1 was unchanged both overall and at individual quantified sites, suggesting that BAP1 does not function as an OGT adaptor protein for all of its direct interactors. The observed dissociation of FOXK1 from OGT upon BAP1 knockout (**Figure 4F**) is consistent with this notion and demonstrates that FOXK1 O-GlcNAcylation does not rely on a stable complex between OGT, BAP1, and FOXK1. Together, these highly localized changes in the network provide experimental support for the adaptor protein hypothesis in the case of some substrates. Moreover, they suggest an expanded, more nuanced form of OGT regulation than simply proximity-enhanced glycosylation, highlighting the potential for both positive and negative regulation of O-GlcNAcylation by OGT adaptor proteins. Overall, our quantitative proteomic studies demonstrate that O-GlcNAcylation can be selectively modulated on a small subset of OGT substrates through manipulation of NISE-identified adaptor proteins in the network.

### Defining conserved and tissue-specific O-GlcNAc networks in vivo

Having established methods to identify functional O-GlcNAc networks in cells, we next expanded our approach in vivo (iNISE). CRISPR/Cas9 technology was used to generate a novel mouse model in which FLAG and HA tags were inserted at the C-terminus of the endogenous OGT gene (OGT-FH, **Figure 5A; Figure S5A**). The OGT-FH mouse line exhibited no obvious phenotypes from development through adulthood. Using western blotting, we confirmed that the expression levels of tagged and untagged OGT were equivalent in major organs of OGT-FH and wild-type (WT) adult mice, indicating that addition of the dual tags did not affect OGT expression levels in vivo (**Figure 5B**).

**Figure 5.**
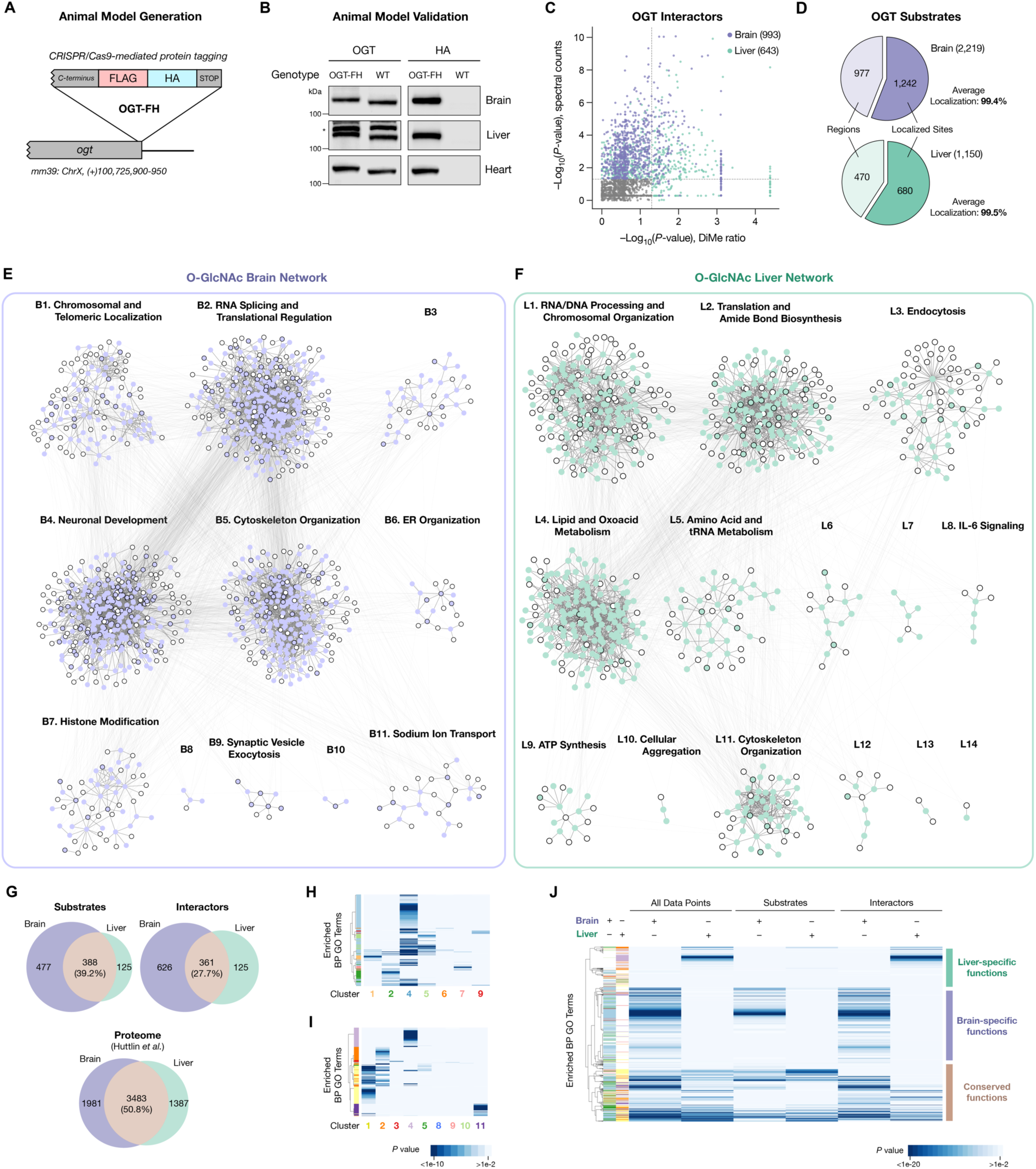
Development and application of O-GlcNAc functional networks in vivo (iNISE). (A) Schematic of genome editing strategy to introduce the FLAG-HA dual tag onto chromosomal OGT. (B) Validation of OGT-FH expression levels equivalent to native OGT across multiple tissues by western blotting. (C) Scatterplot of statistically validated OGT interactors identified from murine brain and liver. (D) Quantification and localization of O-GlcNAc sites identified from murine brain and liver. (E-F) O-GlcNAc functional network from (E) brain or (F) liver with communities annotated by enriched biological process GO terms. (G) Venn diagram of identified OGT interactors and substrates and overall proteomes from murine brain and liver. (H-I) Hierarchical clustering of enriched BP GO terms from initial interactor and substrate datasets and individual GLay communities within the (H) brain or (I) liver networks. (J) Hierarchical cluster of enriched BP GO terms from brain and liver networks highlight the presence of both shared and tissue-specific functions for O-GlcNAcylation.

We next applied iNISE to this OGT-FH mouse model. OGT and its interacting partners were immunoprecipitated from the brain and liver of two-month-old OGT-FH and WT animals. After TAP, proteins were proteolytically digested, and the resulting peptides were isotopically labeled via reductive amination using light (H2CO) or heavy (D2CO) formaldehyde in lieu of SILAC labeling prior to mixing and downstream identification (**Figure S5B**). MS analysis revealed 993 and 643 high-confidence interactors (*n* = 3 per tissue type, **Figure 5C**; **Files S11 and S12**) and 2,219 and 1,150 O-GlcNAc sites (*n* = 2 per tissue type, **Figure 5D**; **Files S13 and S14**) across 810 and 492 proteins (**Files S15 and S16**) from the brain and liver, respectively. A total of 2,785 non-redundant O-GlcNAc sites were definitively identified, which represents the largest number of sites described in a single experiment to date. In this combined dataset, we again saw unique subsets of sites identified only by ETD or EThcD (**Figure S6A**), highlighting the importance of using both fragmentation methods to comprehensively map O-GlcNAc sites.

To organize the myriad of interconnected modification sites into overarching themes, O-GlcNAc functional networks were constructed from these datasets using PPI databases and GLay clustering as described above. Overall, iNISE identified 11 communities in the brain and 14 communities in the liver (**Figures 5E and 5F**; see **Figures S6B and S6C** for the complete annotation of all clusters). Both networks shared clusters containing numerous essential functions, including chromatin organization (Clusters Brain 7 [B7] and Liver 1 [L1]), RNA processing (Clusters B2 and L1), and cytoskeletal organization/dynamics (Clusters B5 and L11). These processes were also shared by HEK293T cells and thus likely represent a core set of OGT functions that are conserved across many cell types and tissues. Notably, each network also contained unique communities with tissue-specific processes, such as neuronal development (Cluster B4) and synaptic vesicle exocytosis (Cluster B9) within the brain network, and endocytosis (Cluster L3), lipid metabolism (Cluster L4), and ATP synthesis (Cluster L9) within the liver network. The overlap between protein substrates in each dataset (39.2%) was relatively similar to the reported overlap between total brain and liver proteomes (50.8%),^100^ whereas the overlap between interactor datasets (27.7%) was more distinct (**Figure 5G**), suggesting that differences in OGT-protein interactions may be primarily responsible for the tissue-specific functions of O-GlcNAcylation. Similar to the HEK293T networks, we found that GO terms were well partitioned into individual clusters in the brain (**Figure 5H**) and liver (**Figure 5I**) networks. Comprehensive hierarchical clustering of all significantly enriched biological process GO terms from both organs revealed both conserved and tissue-specific functions for O-GlcNAcylation (**Figure 5J**). Similarly, we observed both shared and unique elements upon analysis of molecular function, cellular component, complex, pathway, and human phenotype space (**File S17**), providing further evidence of both general and context-specific activities for O-GlcNAcylation.

The in vivo networks provided in-depth information and testable molecular hypotheses regarding O-GlcNAc function and regulation. Using our adaptor protein ranking program, we identified and ranked the top candidate OGT adaptor proteins in the brain and liver (**Figure 6A; Files S18** and **S19**). The RNA-binding protein fox-1 homolog 1 (RBFOX1), a component of the large assembly of splicing regulators (LASR) involved in alternative splicing, autism, and epilepsy,^101^ and the 14-3-3ε protein (YWHAH) were the top-ranked adaptor proteins in the brain and liver, respectively. Interestingly, 14-3-3 proteins bind to protein phosphorylation sites and have also been proposed to function as O-GlcNAc ‘readers’.^102^ Thus, the direct targeting of OGT to 14-3-3 proteins and their phosphorylated binding proteins could provide a powerful means to couple O-GlcNAcylation and phosphorylation in the liver.^103, 104^ Multiple other hits on the OGT adaptor protein lists were annotated with the GO terms “protein binding” and “molecular adaptor activity,” including the postsynaptic density component disks large homolog 1 (DLG1)^105^ and the nuclear importin subunit alpha-5 (KPNA1),^106^ highlighting additional adaptor protein paradigms revealed by iNISE.

**Figure 6.**
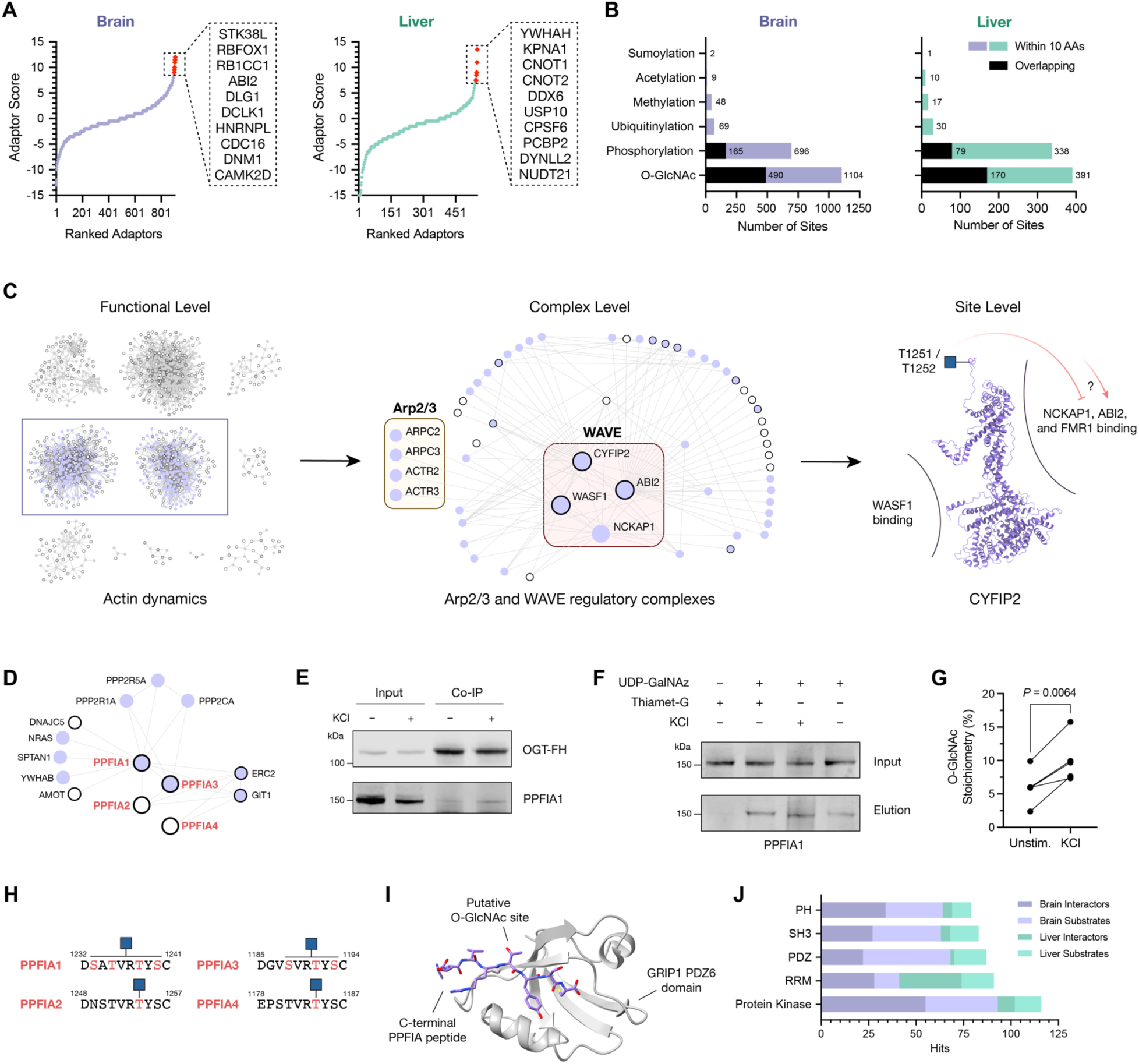
NISE-generated networks reveal new regulatory roles for O-GlcNAcylation in neuronal actin dynamics and synaptic organization. (A) Ranking of potential OGT adaptor proteins within the brain and liver networks. The top ten adaptor proteins for each set are shown. (B) Quantification of PTM sites within the PhosphoSitePlus database that overlap with (black) or are proximal to (color) O-GlcNAc sites within the brain and liver networks. (C) Workflow to identify novel potential O-GlcNAc activities from functional level networks based on cluster annotation, identification of specific OGT-associated protein complexes, and domain- and site-specific information. Structure predicted by AlphaFold. (D) Extracted network of liprin-ɑ (PPFIA, red) proteins and neighboring interactors (solid node) and substrates (outlined node) within the brain network. (E) Validation of the PPFIA1-OGT interaction by co-immunoprecipitation of PPFIA1 with OGT-FH from adult cortical neurons and western blotting. (F-G) Validation and quantification of dynamic PPFIA1 O-GlcNAcylation upon KCl-induced neuronal depolarization or Thiamet-G treatment. O-GlcNAcylated PPFIA1 was detected by chemoenzymatic labeling with UDP-GalNAz and dibenzocyclooctyne-biotin, followed by streptavidin pulldown and western blotting. Paired student’s *t* test. (H-I) Schematic of the highly-conserved putative C-terminal O-GlcNAcylation site within VRTY across PPFIA proteins and its complex with the GRIP1 PDZ6 domain. Structure from PDB 1N7F. (J) Frequency of specific protein domains in OGT interactors and substrates from the brain and liver datasets. PH = pleckstrin homology, SH3 = SRC homology 3, PDZ = PSD-95/Dlg1/ZO-1 homology, RRM = RNA recognition motif.

Incorporation of external existing PTM datasets uncovered a wealth of potential PTM crosstalk within the brain and liver proteomes (**Figure 6B, Files S20 and S21**), with 1,438 PTM sites overlapping or adjacent to O-GlcNAc sites within the brain network alone. Moreover, integration of annotated structural domains found in the interactors and substrates provided additional functional information. For example, protein kinase domains were a recurring feature of many brain and liver network nodes, including six sites and regions on Ca^2+^/calmodulin dependent protein kinase II (CAMK2; T306-T311, T320, T325, S327, T328, and S331-T334) near its Ca^2+^/calmodulin binding domain as well as other well-known regulatory O-GlcNAcylation and phosphorylation sites that modulate its kinase activity during hyperglycemia.^107^ We also uncovered an additional modification region on ULK1 (S746-T768) within a central disordered domain containing AMPK and mammalian target of rapamycin (mTOR) phosphorylation sites that control ULK1 kinase activity and downstream induction of autophagy.^108^ This observation reinforces the general principle that O-GlcNAcylation and phosphorylation play critical, complementary roles in cellular signaling and highlights new possible instances of O-GlcNAc mediated regulation of protein kinase function and specificity.^12, 109^ Incorporation of “disease relevant” functional annotations within the network uncovered potential new molecular mechanisms for PTM interplay in neurological disorders and cancer. These include two O-GlcNAc sites on huntingtin (HTT; T1154 and S1158) near two CDK5-mediated phosphorylation sites that regulate polyQ-induced toxicity in neurons,^110^ as well as a site on nuclear receptor coactivator 3 (NCOA3, also known as steroid receptor coactivator 3 or SRC3; S857) near a phosphorylated serine residue that controls transcriptional activity crucial for tumor progression.^111^

### NISE as a discovery platform for O-GlcNAc regulation

The iNISE-generated networks connect multiple levels of functional and molecular information to facilitate hypothesis-driven investigations into new regulatory roles for O-GlcNAcylation. For example, actin dynamics was a common function shared in the brain and liver. Although O-GlcNAc has been implicated in cytoskeletal remodeling through the modification of proteins such as actin,^112^ myosin,^113^ cofilin,^114^ and protein phosphatase-1 regulator subunit 12A (MYPT1),^86^ our networks suggested expanded roles for O-GlcNAcylation in cytoskeletal reorganization (**Figure 6C**). As several established protein complexes perform actin processing, we first examined the relevant functional communities B4 (nervous system development, cell signaling) and B5 (actin cytoskeleton organization) for known actin machinery using the embedded CORUM complex datasets. Here, we identified multiple key actin-binding proteins and effectors, including the Arp2/3 and WAVE regulatory complexes (WRC), which act together as an actin nucleation site to induce filament branching.^115^ To uncover potential molecular mechanisms of action, we turned to our site-specific O-GlcNAcylation datasets, where we found modification sites on core and accessory members of these complexes, including WRC members Wiskott Aldrich syndrome protein family member 1 (WASF1), cytoplasmic FMR1-interacting protein 2 (CYFIP2), and Abl interactor 2 (ABI2). Query of the public datasets regarding protein structure and function integrated within iNISE then revealed functional hypotheses for these O-GlcNAcylation events. For instance, O-GlcNAcylation of CYFIP2 at T1251 and T1252 lies within the regulatory C-terminal domain that mediates its interactions with WRC component Nck associated protein 1 (NCKAP1) and fragile X mental retardation 1 (FMR1), a key neuronal mRNA binding protein associated with numerous cognitive disorders.^116–118^ Thus, CYFIP2 glycosylation may modulate interactions with NCKAP1, FMR1, or other WRC components to alter complex assembly and thereby affect stimulation-dependent actin polymerization at the neuronal membrane, similar to disease-associated somatic mutations of CYFIP2 characterized in human neurodevelopmental disorders.^119^ Interestingly, ABI2 was both an OGT substrate and interactor that was ranked one of the top 10 adaptor proteins according to our algorithm. Thus, ABI2 may mediate OGT activity toward numerous substrates involved in actin remodeling and synapse formation. In addition to modulating PPIs, iNISE also uncovered potential roles for O-GlcNAcylation in regulating cytoskeletal reorganization via PTM crosstalk. For example, we identified potential regulatory O-GlcNAc sites on the actin-binding protein enabled homology (ENAH) near the protein kinase A phosphorylation site S255 within a region important for neuronal growth cone formation.^120^

Global changes in O-GlcNAcylation are known to affect synapse maturation in both pre and postsynaptic neurons.^10, 121–123^ Although O-GlcNAcylation is highly enriched at neuronal synapses,^55^ few studies have characterized the multiple OGT interactors and substrates involved in synaptic structure and function. The GO biological process data embedded within the iNISE brain network identified 144 OGT interactors and 93 OGT substrates annotated with synaptic function (**File S22**). Through this unbiased analysis, iNISE revealed that all members of the liprin α (PPFIA) family of synaptic scaffolding proteins are O-GlcNAcylated. Moreover, liprin-α1 (PPFIA1) and liprin-α3 (PPFIA3) were identified as putative OGT interactors (**Figure 6D**) and were directly linked in the network to several other proteins important for presynaptic function.^124^ Consistent with these findings, PPFIA1 co-immunoprecipitated with OGT from embryonic cortical neurons cultured from OGT-FH mice (**Figure 6E**). We also confirmed that PPFIA1 was O-GlcNAc modified in cortical neurons using our well-established CL assay^46^ (**Figure 6F**). Notably, we found that KCl-mediated depolarization of cortical neurons induced O-GlcNAcylation of PPFIA1 (**Figure 6G**), suggesting that PPFIA1 glycosylation is dynamically modulated by neuronal activity. PPFIA1 glycosylation occurred within a C-terminal peptide conserved across the PPFIA family of proteins (**Figure 6H**). This peptide region is recognized by the PDZ6 domain of glutamate receptor interacting protein 1 (GRIP1) (**Figure 6I**), which is critical for AMPA receptor clustering in dendrites.^125^ Notably, iNISE revealed that PDZ domains are a common feature of many OGT interactors and substrates in the brain (**Figure 6J**; **File S23**). The association of OGT with PDZ domain-containing proteins important for synaptic organization and plasticity^121^ may allow OGT to regulate pre and post-synaptic signaling on multiple substrates simultaneously and provide a molecular basis for the well-known effects of O-GlcNAcylation on long-term potentiation (LTP) and long-term depression (LTD).^10, 121–123^

## DISCUSSION

Understanding the functional consequences of O-GlcNAcylation events on individual proteins has yielded crucial insights into protein regulation and biological processes across different tissues, disease states, and lifespans. Yet, O-GlcNAcylation and other PTMs act simultaneously on multiple targets, generating many context-specific proteoforms that coordinate biological activity on a global scale. Thus, there remains an urgent need for comprehensive and function-driven approaches that report on the full scope of PTMs within cells to solve this major impediment and advance an understanding of PTM signaling. In this manuscript, we expand the existing framework of PTM interrogation to examine and contextualize O-GlcNAcylation across the proteome for a given biological state. Our work to annotate the multiple functions of O-GlcNAc via the networking of interactors and substrates (NISE) provides a broad platform to drive new functional explorations of this PTM. To obtain the required comprehensive datasets, we developed robust chemical tools, analytical methods, and model systems and established a systematic process to integrate biological and context-specific information. These powerful new tools provided the most extensive in vitro and only in vivo OGT interactome to date (**Files S1**, **S11**, and **S12**). Furthermore, the cross-species O-GlcNAcome of 4,549 sites presented herein represents the largest single collection of non-redundant sites generated in a single study (**Files S2**, **S13**, and **S14**). Altogether, this approach allows for the direct comparison of matched biological samples with unprecedented depth and scale, enabling analyses that were previously not possible.

By organizing these datasets into networks and partitioning the data into distinct communities, NISE creates functionally relevant subclusters that reveal the PPI networks and modification sites likely involved in specific processes. We found that this unbiased method identified numerous examples of O-GlcNAc regulation that were previously reported in the literature, including histone modification,^75, 79^ intracellular transport,^72^ transcription and translation,^5, 126^ synaptic organization,^122, 123^ and carbohydrate metabolism,^6^ demonstrating that the approach successfully uncovers critical functions of O-GlcNAc. At the same time, NISE provided protein and site-specific data to guide mechanistic hypotheses and functional validations. For example, NISE revealed novel roles for O-GlcNAcylation in intron splicing, which can be investigated in future studies by interrogating specific network components crucial for exon stabilization, pre-spliceosome assembly, and intron excision. Moreover, our work provides a valuable, integrated resource for understanding diverse O-GlcNAc activities through the incorporation of information regarding other PTMs, protein domains, regions, motifs, and repeat regions into the networks. Thus, NISE delivers a powerful, systems-level, and unbiased approach to discover the central functions and mechanisms underlying O-GlcNAc activity – a long-standing challenge in the field given the large number of O-GlcNAcylated proteins and the complexity of OGT regulation.

The NISE approach also provides important insights into the upstream regulators and downstream effectors of O-GlcNAc. As described above, testable molecular hypotheses are generated to understand the functional outcomes of protein O-GlcNAcylation in the context of specific biological pathways and processes. The platform is also ideally suited for investigating determinants of O-GlcNAcylation such as adaptor proteins. Through the PPIs built into the networks, we successfully predicted potential OGT adaptor proteins, as well as their prospective targets. Extension of our CL approach, along with state-of-the-art quantitative MS reagents and methods, allowed for the rapid quantification of site-specific changes in O-GlcNAcylation caused by deletion of the adaptor protein BAP1. Consistent with the adaptor protein hypothesis, we found that O-GlcNAc site occupancy changed on only a small subset of the O-GlcNAcome, and changing sites were enriched on substrates proximal to the disrupted interactor in our network. Unexpectedly, we observed both increases and decreases in site occupancy across different substrates and even different sites within the same protein. These observations suggest that adaptor proteins may either enhance or block O-GlcNAcylation of specific substrates. Together, our findings offer new, unexplored mechanisms for selectively modulating the O-GlcNAcylation status of specific proteins or sets of proteins within the proteome.

Finally, the iNISE approach using our newly generated, dual-tagged OGT mouse model has enabled the first comprehensive and comparative study of O-GlcNAc functions in complex tissues. The mouse brain and liver-specific networks, along with the network from cultured human embryonic kidney cells, revealed a core, shared subset of O-GlcNAc activities. These functions centered on the regulation of transcription and translation via epigenetic modifications, transcription factors, and intron splicing, underscoring important, conserved roles of O-GlcNAc spanning the central dogma of molecular biology. The numerous roles of O-GlcNAc in cytoskeletal regulation and organelle transport, which were enriched within all NISE-generated networks, further underscore the critical, broad cellular activities of O-GlcNAc. Within these well-studied functions for O-GlcNAc, the iNISE method detected many novel potential functions and mechanisms of protein regulation beyond modification sites previously reported on core cytoskeletal proteins. For example, modulation of the WAVE regulatory complexes by OGT and O-GlcNAc has not been explored to our knowledge. Together, these molecular discoveries enabled by NISE represent new avenues for understanding the control of cellular morphology important for tissue formation, synaptic function, and oncogenesis. The ability of the NISE approach to rapidly identify novel targets of O-GlcNAc activity was further demonstrated through studies of synaptic proteins in the brain network. We discovered a new, dynamically responsive O-GlcNAc site on the synaptic scaffolding protein PPFIA1 and confirmed its stable interaction with OGT. Integration of protein domain databases within the network revealed that the O-GlcNAc site resides in a C-terminal region that binds to a PDZ domain of GRIP1. In addition, we found that PDZ domains are the third most frequent structural domain found in OGT interactors and substrates. Thus, the regulation of PDZ-containing proteins and their binding partners may be a common feature of O-GlcNAcylation during synaptic development, and one that merits broader examination in the future.

In total, NISE provides a strong foundation to interrogate the activity of PTMs both in vitro and in vivo. As enrichment methods exist for other PTMs, our method could be readily generalized to multiple proteoform states. Although the current approach relies upon existing PPI databases, the advent of new, global interactome datasets will undoubtedly improve both node connectivity within the networks and resulting functional annotations. Moreover, other newly generated resources such as cell type-specific expression levels, protein disorder and structural prediction, germline or disease-specific protein mutations, and small molecule ligandability can also be integrated into the networks to further enable new molecular hypotheses. By applying the tools and methods developed herein, we envision that global analyses of upstream PTM regulators and downstream effectors could be rapidly generated for diverse biological contexts across individual tissues, developmental stages, and disease progression. Thus, the global functional annotation of O-GlcNAcylation will accelerate our understanding of protein regulation and the coordination of PTM activities across the proteome.

## ACKNOWLEDGMENTS

We thank S. Pease and staff at the Caltech Genetically Engineered Mouse Services Core for help with generating the OGT-FH mouse line. This project was supported by the National Institutes of Health (RF1AG060540 to L.C.H.-W. and T32GM008042, T32GM007616, and F30AG055314 to J.W.T), UCLA-Caltech Medical Scientist Training Program (J.W.T), National Science Foundation (GRFP DGE-1144469 to M.E.G.), and Department of Defense (NDSEG to E.H.J.).

## AUTHOR CONTRIBUTIONS

Conceptualization: M.E.G., J.W.T., Y.X., and L.C.H.-W.; Methodology: M.E.G., J.W.T., Y.X., and M.J.S.; Software: J.W.T., M.J.S., and P.C.; Investigation: M.E.G., J.W.T., Y.X., R.B.A., E.H.J., Y.K., A.L.S., T.D.K., A.M., and B.L.; Writing – Original Draft: M.E.G., J.W.T., Y.X., and L.C.H.-W.; Writing – Review & Editing: M.E.G., J.W.T., and L.C.H.-W.; Visualization: M.E.G., J.W.T. and L.C.H.-W.; Supervision: M.E.G., S.D.G., and L.C.H.-W.; Funding Acquisition: L.C.H.-W.

## METHODS

### Chemicals and reagents

Unless otherwise indicated, all chemicals and reagents were obtained from MilliporeSigma (Darmstadt, Germany) and used as provided.

### Cell culture conditions

The parental HEK293T cell line was obtained from American Type Culture Collection (Manassas, VA). All cell culture reagents were purchased from Thermo Fisher Scientific (Waltham, MA) unless otherwise indicated. HEK293T cells were cultured in DMEM supplemented with high glucose, GlutaMAX, 10% fetal bovine serum (FBS), and 1x penicillin–streptomycin (P/S) (complete DMEM). To perform quantitative proteomics, cells were cultured for at least six passages in SILAC medium. SILAC medium was produced using SILAC DMEM Flex media supplemented with high glucose, GlutaMAX, 10% dialyzed FBS, 1x P/S and either 0.80 mM L-lysine and 0.40 mM L-arginine (MilliporeSigma; light; K0, R0) or 0.80 mM [U-^13^C_6_,^15^N_2_] L-lysine and 0.40 mM [U-^13^C_6_] L-arginine (Cambridge Isotope Laboratories, Tewksbury, MA; heavy: Lys8, Arg6).

### Lentiviral plasmid construction

All primers were purchased from Integrated DNA Technologies (Coralville, IA), and all molecular biology supplies were purchased from New England Biolabs (Ipswich, MA), unless otherwise indicated. The OGT lentiviral vector was produced by first cloning OGT with C-terminal FLAG and HA tags into the pCMV6 entry vector (Origene, Rockville, MD) using the NEBuilder HiFi DNA assembly and Q5 site-directed mutagenesis kits according to the manufacturer’s protocols. Both the tagged form of OGT (OGT-FH) and pTRIPZ vector were linearized by PCR amplification using the Q5 Hot Start High-Fidelity 2x master mix. Fragments were gel purified using the Zymoclean gel DNA recovery kit (Zymo Research, Tustin, CA) and assembled with the NEBuilder HiFi DNA assembly kit according to the manufacturer’s protocol. The ligated plasmid was transformed into Stable competent cells (New England Biolabs), and individual clones were picked and sequenced (Laragen, Culver City, CA) to ensure proper incorporation and no recombination. Bacterial cultures were stored as 50% glycerol stocks at -80 °C, and 100-mL maxiprep cultures of the plasmid-expressing bacteria were grown and purified by ZymoPure plasmid kits (Zymo Research). Each purification of the pTRIPZ-OGT-FH plasmid was resolved by 1% agarose gel to ensure no recombination occurred.

### Cell line generation

Transfection and cell culture reagents were purchased from Thermo Fisher Scientific unless otherwise indicated. The lentiviral packaging plasmids pMD2.G and psPAX2 (Didier Trono; 12259, 12260) were obtained from Addgene (Cambridge, MA). Four 15-cm culture plates containing HEK293T cells at 80% confluency were each transfected with 8 μg pTRIPZ-OGT-FH, 6 μg psPAX2, and 2 μg pMD2.G using Lipofectamine 3000 at a 2:1 lipid/DNA ratio in OptiMEM according to the manufacturer’s protocol. The medium was replaced at 6 h post-transfection with complete DMEM containing 2% FBS. Medium was harvested at 48 h and 72 h post-transfection, and lentiviral particles were concentrated using 15-mL 100 kDa MWCO Amicon concentrator tubes (MilliporeSigma). Lentiviral concentrate was used immediately. To a six-well plate containing HEK293T cells at 80% confluency in complete DMEM with 8 μg mL^-1^ hexadimethrine bromide, 0-50 μL lentiviral concentrate was added. Medium was replaced with complete DMEM after 24 h. After 48 h, cells were passaged 1:4 into new six-well plates with complete DMEM containing 2 μg mL^-1^ puromycin (Thermo Fisher Scientific). After selection for two weeks, surviving cells were then split for clonal dilution in 96-well plates. After 2-3 weeks, single clones were isolated and expanded. Clones were then tested for induction of OGT-FH expression by incubating with complete DMEM containing 0.1 μg mL^-1^ doxycycline hyclate for 24 h followed by western blotting. A single clone for the HEK293T-iOGT-FH cell line was then chosen and used for subsequent experiments.

### Mouse line generation and genotyping

All mouse procedures were performed in accordance with protocols approved and guidelines set by the Caltech Institutional Animal Care and Use Committee (Protocol IA21-1435). All mice were obtained from Jackson Laboratory (Bar Harbor, ME) or Charles River (Wilmington, MA). The OGT-FH mouse model was maintained on a C57BL/6J background. sgRNA candidates were designed using the CRISPR design (www.genome-engineering.org) and CHOPCHOP programs (chopchop.cbu.uib.no). Genomic DNA was isolated from the tail tip of a wild-type C57BL/6J mouse using the DNEasy Blood and Tissue Kit (Qiagen, Hilden, Germany). The genomic C-terminal region of OGT was amplified using the Q5 Hot Start High Fidelity 2x master mix (New England Biolabs). The pCAG-EGxxFP plasmid (Addgene, Masahito Ikawa; 50716) was cut with BamHI and SalI, and the construct was assembled using the NEBuilder HiFi Assembly kit. sgRNA candidates were cloned into the pX330 plasmid (Addgene, Feng Zhang; 42230) according to the published protocol. The two plasmids were then transfected into HEK293T cells at 80% confluency in a six-well plate using Lipofectamine 3000 at a 2:1 lipid/DNA ratio according to the manufacturer’s protocol. After 48 h, cells were observed using an LSM 710 confocal microscope (Carl Zeiss AG, Oberkochen, Germany) for GFP^+^ cells, indicating active sgRNA. The most active sgRNA found within 20 bp of the OGT stop codon was chosen for *in vivo* genomic editing (5’-CCTGAATAAAGACTGCGCAC-3’).

The sgRNA was amplified from the pX330 plasmid and then transcribed and purified using the MEGAshortscript T7 transcription and clean-up kits (Thermo Fisher Scientific), respectively. For homology-directed recombination, a single-stranded oligodeoxynucleotide (ssODN) was synthesized as an Ultrimer oligonucleotide (IDT) containing an MluI cut site, FLAG tag, BamHI cut site, HA tag, and stop codon flanked on either side by 60 nucleotides homologous to the genomic region surrounding the insertion site. The protospacer adjacent motif was mutated to prevent further nuclease activity after homology-directed recombination.

Zygotes from C57BL/6N mice (Charles River) were produced, collected, cultured, and implanted as previously described.^127^ Microinjection of embryos was performed using an inverted microscope (Carl Zeiss AG) equipped with a micromanipulator (Leica Microsystems, Wetzlar, Germany), CellTram (Eppendorf, Hamburg, Germany), and FemtoJet (Eppendorf). Injections were carried out as previously described.^128^ Solutions containing 2.5 ng μL^-1^ sgRNA, 10 ng μL^-1^ ssODN, and 5 ng μL^-1^ Cas9 mRNA (System Biosciences, Palo Alto, CA) in 10 mM Tris-HCl pH 8.0 and 0.1 mM EDTA were used for injection into the pronucleus of fertilized mouse zygotes. Following injection, the zygotes were cultured to the two-cell stage and implanted into pseudopregnant foster mothers at 0.5 days post coitum (up to 30 two-cell embryos per recipient). Approximately 19.5 days after implantation, the pups were delivered, and 3 weeks after birth the pups were tailed and separated by gender. Offspring were genotyped by restriction enzyme digestion using BamHI after purification of the genomic DNA from tail tips using the DNEasy Blood and Tissue kit and PCR amplification of the genomic C-terminal region of OGT. A single heterozygous female was obtained containing the correct on-target inserted sequence. The line was then backcrossed for at least six generations with the C57BL/6J strain prior to experiments. Subsequent homozygous mice (OGT-FH mice) obtained from this founder exhibited no abnormalities in growth, behavior, or breeding and were indistinguishable from wild-type and heterozygous mice.

### siRNA knockdown

The BAP1 RNA knockdown was performed with a set of 3 unique 27-mer siRNA duplexes targeting BAP1 (Origene, SR305435) using siTrans 1.0 (Origene). Briefly, cells in six-well plates at 70% confluency were transfected with 20 nmol siRNA mixture according to the manufacturer’s protocol. Cells were allowed to grow for three days prior to harvesting as described for cell lysates above. Knockdown was validated by western blotting. Co-immunoprecipitation experiments were carried out in the same manner as described for TAP-MS sample preparation.

### CRISPR/Cas9-mediated knockout of BAP1

BAP1 CRISPR/Cas9 KO Plasmid (h) (sc-400232), BAP1 HDR Plasmid (h) (sc-400232-HDR), and Control CRISPR/Cas9 Plasmid (sc-418922) were purchased from Santa Cruz Biotechnology (Dallas, TX) and prepared per the manufacturer’s instructions. HEK293T cells at 60-70% confluency (maintained as described above) in a 12-well tissue culture plate were then transfected with 1 ug of BAP1 KO/HDR or control plasmid using Lipofectamine 3000 (Thermo Fisher Scientific) at a 2:1 lipid/DNA ratio in OptiMEM according to the manufacturer’s protocol. Once confluent, the cells were split 1:1 into media containing 10 μg mL^-1^ puromycin (Thermo Fisher Scientific) or normal media for the control. Cells were then selected for 2 weeks replacing the media or splitting as necessary. BAP1 knockout was confirmed by western blotting. KO and WT cells were grown in three 10-cm plates, lysed, and subjected to chemoenzymatic labeling and enrichment as described below. Lysate (100 μg) from each condition was also saved for quantitative proteomics.

### Cell lysate preparation

To prepare HEK293T cell lysate for immunoprecipitation or western blotting, two 15-cm plates of heavy-labeled HEK293T-iOGT-FH cells at 60% confluency were treated with freshly made 0.04 μg mL^-1^ doxycycline hyclate for 24 h. As a control, light-labeled HEK293T-iOGT-FH cells were left untreated. The cells were collected with a scraper and homogenized on ice in 10 mL of 50 mM Tris-HCl pH 7.4, 150 mM NaCl, 0.5% NP-40, and 1x cOmplete EDTA-free protease inhibitor cocktail (MilliporeSigma) using 30-40 strokes with a 20-mL Potter-Elvehjem tissue grinder (Wheaton, Millville, NJ) followed by centrifugation at 15,000 g for 10 min at 4 °C. The protein concentration of the clarified supernatant was measured using the Pierce BCA Protein Assay Kit (Thermo Fisher Scientific).

To prepare HEK293T cell lysate for chemoenzymatic labeling, two 15-cm plates of HEK293T-iOGT-FH or three 10-cm plates of HEK293T-BAP1KO cells at 60% confluency were lysed in 5 mL of 1% SDS, 50 mM Tris-HCl pH 7.4, 150 mM NaCl, 1x cOmplete EDTA-free protease inhibitor cocktail, and 10 μM Thiamet-G and probe sonicated to shear DNA. The protein concentration was measured as described above.

### Affinity purification from 293T cells

For each replicate, approximately 15 mg of cell lysate from heavy- and light-labeled cells were subjected to immunoprecipitation with 200 μL settled volume of ANTI-FLAG M2 Affinity Agarose Gel (MilliporeSigma) overnight at 4 °C. In all cases, 1% of the sample (approximately 150 μg) was saved as input for total protein staining and western blotting. The beads were washed 3 x 5 min in wash buffer (50 mM Tris-HCl pH 7.4, 150 mM NaCl, and 0.1% NP-40) and eluted with 150 μg mL^-1^ of 3xFLAG peptide (MilliporeSigma) in wash buffer, followed by immunoprecipitation with 100 μL settled volume of anti-HA agarose beads (Thermo Fisher Scientific) for 6 h at 4 °C. The beads were washed with wash buffer and eluted with 3 M NaSCN. The eluates were acetone precipitated, re-dissolved in 1x SDS loading buffer (50 mM Tris-Cl pH 6.8, 100 mM dithiothreitol (DTT), 2% SDS, 0.1% bromophenol blue, 10% glycerol), and subjected to in-gel digestion.

The eluates from heavy- and light-labeled samples were mixed immediately before loading onto a NuPAGE 4-12% Bis-Tris protein gel (Thermo Fisher Scientific). The gel was stained and visualized using Imperial protein staining reagent (Thermo Fisher Scientific) and destained in doubly deionized H_2_O (ddH_2_O). The gel lane was cut into 30-40 gel slices. Each gel piece was reduced with 10 mM DTT for 1 h at 37 °C and then alkylated with 50 mM iodoacetamide for 45 min at room temperature protected from light. The slides were dried and incubated with 20 ng μL^-1^ trypsin protease (Pierce MS Grade, Thermo Fisher Scientific) solution in 100 mM NH_4_HCO_3_ pH 8.0 and 1 mM CaCl_2_ overnight at 37 °C. Peptides were recovered using sequential washes with 100 mM NH_4_HCO_3_, 1:1 v/v 100 mM NH_4_HCO_3_/CH_3_CN, and 5% formic acid. The recovered peptides were dried and desalted with a C18 ZipTip pipette tip (MilliporeSigma). Finally, the desalted peptides were resuspended in 0.2% formic acid for LC-MS/MS analysis.

### Affinity purification from brain and liver tissue

To prepare brain and liver tissue lysates, 2-month-old OGT-FH and WT mouse forebrains and livers were freshly dissected and rinsed with ice cold phosphate-buffered saline (PBS). One brain hemisphere or liver lobe was used per experiment. Lysate from each of OGT-FH and WT organs was harvested and homogenized on ice in 10 mL of 50 mM Tris-HCl pH 7.4, 150 mM NaCl, 0.5% NP-40, and 1x cOmplete EDTA-free protease inhibitor cocktail using 30-40 strokes with a Potter-Elvehjem homogenizer followed by centrifugation at 15,000 g for 10 min at 4 °C. Lysates (15 mg) from each mouse were subjected to immunoprecipitation using the same procedure outlined above, followed by filter assisted sample preparation^129^ and dimethyl labeling.^130^

Briefly, the bead eluates were loaded onto 0.5-mL 10 kDa MWCO Amicon Ultra centrifugal filters (MilliporeSigma) and buffer exchanged into 100 mM TEAB pH 8.5. The samples were then reduced with 10 mM DTT for 1 h at 37 °C and alkylated with 50 mM iodoacetamide for 45 min at room temperature protected from light. The proteins were digested with 50 ng μL^-1^ trypsin in 100 mM TEAB at 37 °C overnight. The resulting peptides were recovered by washing the centrifugal filter with 100 mM TEAB three times for a final volume of 100 μL. To each peptide mixture, 4 μL of 4% v/v CH_2_O (WT) or CD_2_O (OGT-FH) was added followed by 4 μL of 0.6 M NaBH_3_CN. Samples were then rotated end-over-end for 1 h at room temperature and quenched with 16 μL of 1% v/v NH_3_ followed by 8 μL of formic acid. The differentially labeled samples from OGT-FH and WT mice were mixed and desalted by an Agilent 1100 Series HPLC system with a MicroTrap Cartridge (Michrom, 1 x 8 mm I.D., 5.0 μL bed volume, 20 μg capacity, 1-1000 μL sample volume). Solvent A consisted of 99.8% H_2_O and 0.2% formic acid and solvent B consisted of 99.8% CH_3_CN, and 0.2% formic acid. The gradient was as follows: 0% solvent B (10 min), 0-85% B (2 min), 85% B (5 min), 85-90% B (1 min), 90% B (5 min), 90-100% B (1 min), and 100% B (6 min). Eluted peptide fractions were collected in 1-mL fractions, lyophilized, and resuspended in 0.2% formic acid for LC-MS/MS analysis.

### Chemical synthesis of biotin-Dmpt-alkyne 1

Reactions were performed in flame-dried glassware under Ar using freshly dried solvents passed through an activated alumina column under Ar. Preparative HPLC was conducted using a 1260 Infinity II LC system (Agilent) with a custom Zorbax Eclipse XDB-C18 PrepHT column (Agilent, 21.2 x 250 mm, 5 μm) with a linear gradient of 5-50% CH3CN with 0.1% formic acid over 70-85 min with a flow rate of 6 mL min^-1^. ^1^H and ^13^C NMR experiments were recorded on a Bruker 400 spectrometer at the Caltech NMR Facility. Spectra are reported in parts per million (δ) relative to CDCl3 (7.26 ppm). Data are reported as chemical shift (ppm), multiplicity (s = singlet, d = doublet, t = triplet, q = quartet, p = pentet, m = multiplet, b = broad), coupling constant (Hz), and integration. Mass spectra were obtained using a Waters LCT Premier XE electrospray time of flight mass spectrometer at the Caltech Multi-User Mass Spectrometry Laboratory.

#### 5-(3-(2-(2-(2-(2-(*N*-biotinyl)aminoethoxy)ethoxy)ethoxy)ethoxy)-1-hydroxypropylidene) 1,3-dimethylpyrimidine-2,4,6-trione (Biotin-Dmpt-OH, 2)

Biotin-PEG4-CO2H (BroadPharm, 400 mg, 0.814 mmol) and *N,N’*-dimethylbarbituric acid (165 mg, 1.06 mmol, 1.3 eq) were suspended in 10 mL of dry dichloromethane (DCM) under Ar and cooled to 0 °C. Separately, 1-(3-dimethylaminopropyl)-3-ethylcarbodiimide hydrochloride (EDC·HCl, 203 mg, 1.06 mmol, 1.3 eq), DMAP (9.9 mg, 0.0810 mmol, 0.1 eq), and TEA (170 μL, 1.22 mmol, 1.5 eq) were dissolved in 5 mL of dry DCM. The coupling solution was added to the suspension dropwise. The reaction was allowed to warm to room temperature and stirred overnight. The solution was then concentrated to a yellow oil and redissolved in a minimal volume of 5% aqueous CH_3_CN. The product was then purified by preparative HPLC. Fractions were combined, concentrated, and lyophilized to afford the product as a pale yellow powder (469 mg, 92% yield). ^1^H NMR (500 MHz, CDCl_3_): δ 7.33 (bs, 0.2H, O*H*), 6.73 (bt, *J* = 5.4 Hz, 0.8H, O*H*), 6.28 (s, 1H, N*H*), 5.43 (s, 1H, N*H*), 4.51 (t, *J* = 6.5 Hz, 1H, biotin-C*H*), 4.32 (dd, *J* = 7.6, 4.5 Hz, 1H, biotin-C*H*), 3.86 (t, *J* = 6.3 Hz, 2H, OC*H*_2_), 3.68-3.59 (m, 12H, OC*H*_2_), 3.55 (t, *J* = 5.1 Hz, 2H, C*H*_2_), 3.48-3.27 (m, 10H, OC*H*_2_, C*H*_3_), 3.15 (dt, *J* = 12.0, 6.0 Hz, 1H, biotin-C*H*), 2.91 (dd, *J* = 12.7, 4.9 Hz, 1H, biotin-C*H*), 2.73 (d, *J* = 12.7 Hz, 1H, biotin-C*H*), 2.22 (td, *J* = 7.1, 3.4 Hz, 2H, biotin-C*H*_2_), 1.82-1.57 (m, 4H, biotin C*H*_2_), 1.44 (p, *J* = 7.5 Hz, 2H, biotin-C*H*_2_). ^13^C NMR (100 MHz, CDCl_3_): 196.88, 173.41, 169.85, 164.02, 150.45, 96.02, 77.36, 70.68, 70.57, 70.53, 70.34, 70.22, 70.08, 66.91, 61.91, 60.33, 55.55, 40.68, 39.29, 37.26, 35.96, 28.21, 25.65. ESI-HRMS (*m/z*): [M+H]^+^ calc’d for C_27_H_44_N-_5_O_10_S^+^: 630.2803, found: 630.2791.

#### 5-(3-(2-(2-(2-(2-(*N*-biotinyl)aminoethoxy)ethoxy)ethoxy)ethoxy)-1-(*N* propargyl)aminopropylidene)-1,3-dimethylpyrimidine-2,4,6-trione (Biotin-Dmpt-alkyne, 1)

Compound **2** (50.0 mg, 0.0794 mmol) was suspended in propargylamine (2.00 mL, 31.2 mmol) and stirred at 50 °C for 4 h. The reaction was then concentrated, and the residue was diluted in DCM. The solution was extracted with H_2_O, and the aqueous layer was back-extracted twice with DCM. The organic layers were combined, dried with MgSO_4_, and concentrated. The residue was redissolved in a minimal volume of 5% aqueous CH_3_CN and then purified by preparative HPLC. Fractions were combined, concentrated, and lyophilized to afford the product as a pale yellow powder (14.8 mg, 28% yield). ^1^H NMR (400 MHz, CDCl3): δ 12.87 (bt, *J* = 5.5 Hz, 1H, N*H*), 6.64 (bt, *J* = 5.6 Hz, 1H, N*H*), 6.20 (s, 1H, N*H*), 5.28 (s, 1H, N*H*), 4.50 (ddt, *J* = 7.5, 5.0, 1.2 Hz, 1H, biotin-C*H*), 4.45 (dd, *J* = 5.5, 2.6 Hz, 2H, NC*H*_2_), 4.31 (ddd, *J* = 7.8, 4.6, 1.4 Hz, 1H, biotin-C*H*), 3.86 (t, *J* = 5.5 Hz, 2H, OC*H*_2_), 3.67-3.53 (m, 14H, OC*H*_2_), 3.51-3.40 (m, 4H, OC*H*_2_), 3.33 (s, 3H, NC*H*_3_), 3.29 (s, 3H, NC*H*_3_), 3.14 (td, *J* = 7.4, 4.6 Hz, 1H, biotin-C*H*), 2.90 (dd, *J* = 12.8, 4.9 Hz, 1H, biotin-C*H*), 2.74 (d, *J* = 12.8 Hz, 1H, biotin-C*H*), 2.41 (t, *J* = 2.6 Hz, 1H, CC*H*), 2.22 (td, *J* = 7.3, 2.2 Hz, 2H, biotin-C*H*_2_), 1.78-1.58 (m, 4H, biotin-C*H*_2_), 1.44 (q, *J* = 7.6 Hz, 2H, biotin-C*H*_2_). ^13^C NMR (100 MHz, CDCl_3_): δ 175.65, 173.30, 166.84, 163.79, 162.63, 151.35, 90.31, 77.86, 77.36, 73.87, 70.64, 70.62, 70.56, 70.52, 70.25, 70.12, 69.91, 61.88, 60.28, 55.56, 40.68, 39.28, 36.02, 33.97, 31.14, 28.25, 28.22, 28.14, 27.91, 25.67. ESI-HRMS (*m/z*): [M+H]^+^ calc’d for C_30_H_47_N-_6_O_9_S^+^: 667.3120, found: 667.3133.

### Chemoenzymatic labeling and enrichment for O-GlcNAcomics

HEK293T-iOGT-FH cell lysate, brain, liver, or WT/BAP1 KO 293T cell lysate (5 mg per condition in all cases) was diluted to 2.5 mg mL^-1^ with 1% SDS and 20 mM 4-(2-hydroxyethyl)-1 piperazineethanesulfonic acid (HEPES) pH 7.6. Samples were then reduced using 20 mM DTT in 20 mM HEPES pH 7.6 for 45 min at 60 °C, cooled to room temperature, and alkylated with 100 mM iodoacetamide for 45 min at room temperature protected from light. Protein was then precipitated by the addition of 3 volumes of CH3OH, 1 volume of CHCl3, and 2.25 volumes of H2O. The precipitated protein was pelleted by centrifugation at 7,068 x *g* for 35 min at 4 °C. Solvents were removed and the pellet washed twice with 3 volumes of CH3OH before redissolving at 5 mg mL^-1^ with 1 mL of 1% SDS and 20 mM HEPES pH 7.9 at 95 °C for 5 min. Samples were then diluted with 2 mL of 2.5x GalT labeling buffer (50 mM HEPES pH 7.9, 125 mM NaCl, 5% IGEPAL CA-630), 1.2 mL of ddH2O, and 275 μL of 100 mM MnCl2. The mixture was chilled on ice for 5 min, and then 250 μL of 0.1 mM UDP-GalNAz^131^ in 10 mM HEPES pH 7.9, 250 μL of 2 mg mL^-1^ Y289L GalT,^46^ 50 μL of 500 kU mL^-1^ PNGase F (New England Biolabs, 5 U μg^-1^ of protein), and 62.5 μL of Lambda Protein Phosphatase (New England Biolabs, 5 U μg^-1^ of protein) were added with gentle mixing after each addition. Samples were rotated end-over-end overnight at 4 °C and were then precipitated by the CH_3_OH/CHCl_3_/H_2_O method described above. The protein pellet was redissolved at 5 mg mL^-1^ in 1 mL of 1% SDS and 20 mM HEPES pH 7.6 at 95 °C for 5 min. The resulting solution was diluted with 2.85 mL of ddH_2_O and 0.15 mL of 200 mM HEPES pH 7.6. In a separate tube, a 5x CuAAC mix was prepared by sequential addition of 885 μL of ddH_2_O, 5 μL of 100 mM BTTAA (Click Chemistry Tools, Scottsdale, AZ) in DMSO, 100 μL of 50 mM CuSO_4_ (freshly prepared), and 10 μL of 50 mM **1** in DMSO. The 5x mix was then added directly to the protein sample and mixed gently before adding 100 μL of 100 mM TCEP (freshly prepared). The CuAAC reaction was allowed to proceed with end-over-end rotation for 1 h at room temperature before the proteins were precipitated as described above. The pellet was resuspended in 5 mL of 50 mM HEPES pH 7.6 and 10 mM EDTA to chelate residual Cu^2+^. Then, 200 μg of trypsin protease (1:25 w/w) was added, and the sample was rotated end-over-end for 20 h at 37 °C. Two 15-mL 10 kDa MWCO Amicon concentrator tubes were rinsed with 2 x 5 mL of 50% CH3OH and 2 x 5 mL of ddH2O by centrifugation at 4,000 x *g* for 5 min. Tryptic solutions were then centrifuged in the rinsed concentrator tubes at 4,000 x *g* for 20 min. The remaining residue was rinsed 2 x 2.5 mL of H2O by centrifugation, and the flowthrough fractions were combined and diluted two-fold with PBS. In a separate tube, 500 μL of a 50% slurry of high capacity Neutravidin agarose (Thermo Fisher Scientific) was washed twice with 0.5 mL of PBS. The washed agarose resin was then added to the sample and rotated end-over-end for 1 h at room temperature. The resin was pelleted by centrifugation at 500 x *g* for 5 min and then transferred to a 900-μL spin filter (Thermo Fisher Scientific) pre-rinsed twice with 1:1 CH3OH/ddH2O. Beads were washed with the following solutions: 5 x 0.5 mL of PBS, 5 x 0.5 mL of 1 M NaCl in PBS, and 5 x 0.5 mL of PBS. Captured peptides were then eluted twice with 0.75 mL of 2% *v*/*v* NH2OH with end-over-end rotation for 1.5 h at 37 °C. Elution fractions were combined and concentrated to dryness with vacuum centrifugation before desalting as described for affinity purified peptides. For each pair of quantitative samples, dried peptides were resuspended in 50 μL of 100 mM TEAB pH 8.5 and labeled using two tags from the TMT10plex Isobaric Labeling Kit (Thermo Fisher Scientific) per the manufacturer’s instructions. Labeled peptides were then combined, desalted and dried as before, and resuspended in 10 μL of 0.2% formic acid for MS analysis.

### Preparation of peptides for quantitative proteomics experiments

WT and BAP1 KO lysates (100 μg each, n = 3 for each) were prepared separately as described above and precipitated using the CH_3_OH/CHCl_3_/H_2_O method as described above except that the samples were centrifuged at 21,130 x g for 5 min to pellet proteins. 100 μl of 100 mM TEAB pH 8.5 was added to the pellets, followed by 3 μg of trypsin protease (1:33 w/w enzyme:protein). The samples were digested for 20 h at 37 °C and then labeled with the TMTsixplex Isobaric Labeling Kit (Thermo Fisher Scientific) per the manufacturer’s instructions. After labeling, peptides from all groups were combined and concentrated to dryness by vacuum centrifuge. Peptides were then resuspended in 100 μL of 10 mM NH_4_OH and subjected to offline high pH, reversed-phase fractionation^132^ using an Agilent Extend-C18 (2.1 x 150 mm, 5 μm) on an Agilent 1100 HPLC operating at 0.2 mL min^-1^. Solvent A consisted of 10 mM NH_4_OH, and solvent B consisted of 10 mM NH_4_OH in 90% CH_3_CN. The combined peptide sample (600 μg) was injected onto the column and separated per the following gradient: 1% solvent B (4 min), 1-30% B (50 min), 30-60% B (4 min), 60-70% B (2 min), and 70-90% B (5 min). 64 1-min fractions were collected in 96 deep-well plates, concatenated to 16 samples, and dried by vacuum centrifugation. Finally, each fraction was resuspended in 10 µL of 0.2% formic acid by bath sonication for MS analysis.

### LC-MS/MS analysis

#### Interactome samples after affinity purification from HEK293T cells

Affinity purified, in-gel digested samples from HEK293T cells were subjected to LC-MS/MS analysis on a nanoflow LC system, EASY-nLC II (Thermo Fisher Scientific), coupled to a LTQ Orbitrap Elite mass spectrometer (Thermo Fisher Scientific). For the EASY-nLC II system, solvent A consisted of 97.8% H_2_O, 2% CH_3_CN, and 0.2% formic acid, and solvent B consisted of 19.8% H_2_O, 80% CH_3_CN, and 0.2% formic acid. Samples were directly loaded onto a 16-cm analytical HPLC column (50 µm I.D.) packed in-house with ReproSil-Pur C18AQ 3 µm resin (120 Å pore size, Dr. Maisch, Ammerbuch, Germany). The column was heated to 60 °C during separation. The peptides were separated with a 60-min gradient at a flow rate of 220 nL min^-1^ with the following gradient: 2-30% solvent B (60 min), 30-100% B (1 min), and 100% B (9 min). Eluted peptides were then ionized using a Nanospray Flex ion source (Thermo Fisher Scientific) and introduced into the mass spectrometer. The LTQ Orbitrap Elite was operated in a data-dependent mode, automatically alternating between a full-scan (400-1600 *m/z*, 120K resolution) in the Orbitrap and subsequent MS^2^ scans of the 20 most abundant peaks in the linear ion trap (Top20 method).

#### Interactome samples after affinity purification from brain and liver tissue

Affinity purified samples from brain tissue were subjected to LC-MS/MS analysis on a nanoflow LC system, EASY-nLC 1200 (Thermo Fisher Scientific), coupled to a Q Exactive HF Orbitrap mass spectrometer (Thermo Fisher Scientific) equipped with a Nanospray Flex ion source. Samples were directly loaded onto a 20-cm PicoFrit column (50 µm I.D., New Objective, Woburn, MA) packed in house with ReproSil-Pur C18AQ 1.9 µm resin (120 Å pore size, Dr. Maisch, Ammerbuch, Germany). The column was heated to 60 °C during separation. The peptides were separated with a 120-min gradient at a flow rate of 220 nL min^-1^ with the following gradient: 2-6% solvent B (7.5 min), 6-25% B (82.5 min), and 25-40% B (30 min) and to 100% B (9 min). Solvents A and B were the same as described above. The Q Exactive HF Orbitrap was operated in data dependent mode. Full scan resolution was set to 60,000 at 200 *m/z*. Full scan target was 3 × 10^6^ with a maximum injection time of 15 ms. Mass range was set to 300-1650 *m/z*. For data dependent MS^2^ scans the loop count was 12, target value was set at 1 × 10^5^, and intensity threshold was kept at 1 × 10^5^. Isolation width was set at 1.2 *m/z* and a fixed first mass of 100 was used. Normalized collision energy was set at 28%. Peptide match was set to off, and isotope exclusion was on.

#### O-GlcNAcomics samples

Chemoenzymatically labeled, enriched, and digested samples were analyzed on a nanoflow LC system, EASY-nLC 1000 (Thermo Fisher Scientific), coupled to an Orbitrap Fusion Tribrid mass spectrometer (Thermo Fisher Scientific) equipped with a Nanospray Flex ion source. Sample loading and LC separation was identical to the Q Exactive HF Orbitrap LC method outlined above. Full spectra were acquired over 350-1800 *m/z* in the Orbitrap (120 K resolution at 200 *m/z*); automatic gain control (AGC) was set to accumulate 50,000 ions, with a maximum injection time of 50 ms. Data-dependent MS^2^ analysis was performed using a top speed approach (cycle time of 5 seconds) with multiple fragmentation methods (see below). Dynamic exclusion was set to exclude features after 1 time for 15 seconds with exclude isotopes turned on. The normalized collision energy was optimized at 28% for high collision dissociation (HCD) fragmentation. The intensity threshold for fragmentation was set to 25,000. HCD fragmentation spectra were collected in the Orbitrap operating at 30K resolution at 200 *m/z*. ETD/EThcD fragmentation was then performed for precursor ions whose HCD spectra contained a fragment of mass 300.1303 *m/z* (15 ppm tolerance, within the top 30 ions). AGC was set to 50,000 with a maximum injection time set at 500 ms for the Orbitrap operating at 30K resolution. ETD reaction time was charge-dependent and supplemental activation (SA) energy was set to 20%.

#### Quantitative O-GlcNAcomics samples

Quantitative O-GlcNAcomics samples were analyzed on a nanoflow LC system, EASY nLC 1000, coupled to an Orbitrap Fusion Tribrid mass spectrometer equipped with a Nanospray Flex ion source. ∼4 μg of peptides per sample were loaded onto a 25-cm Aurora column (75 µm I.D.) packed in-house with 1.6 µm C18 resin (Ion Opticks, Parkville, Victoria, Australia). Solvents A and B were the same as described above for interactome samples. Peptides were separated over 135 min at a flow rate of 350 nL min^-1^ with the following gradient: 2-6% solvent B (7.5 min), 6-25% B (82.5 min), 25-40% B (30 min), 40-100% B (1 min), and 100% B (14 min). MS1 spectra were acquired at 120K resolution with a scan range from 350-1800 *m/z*. AGC was set to accumulate 50,000 ions, with a maximum injection time of 50 ms. Data-dependent MS^2^ analysis, either top speed (5 seconds) or top 10, was then performed in which features were filtered for monoisotopic peaks with a charge state of 3-8 and a minimum intensity of 25,000, with dynamic exclusion set to exclude features after 1 time for 15 seconds with a 10 ppm mass tolerance and exclude isotopes turned on. HCD fragmentation was performed with normalized collision energy of 28% after quadrupole isolation of features using an isolation window of 1.6 *m/z*, an AGC target of 1 x 10^5^, and a maximum injection time of 100 ms. MS^2^ scans were then acquired at 30K resolution in Centroid mode with the scan range set to automatic. Detection of at least one fragment with 732.3726, 547.3037, or 529.2931 *m/z* in the top 30 ions triggered three additional scans: (1) SPS-MS^3^ in the Orbitrap on the top 5 SPS precursors, an isolation window of 1.3 *m/z*, 65% normalized collision energy, a resolution of 50K, an ACG target of 50,000, an automatic scan range, and a maximum injection time of 250 ms. Precursor selection range, ion exclusion, and isobaric tag loss exclusion were all employed; (2) ETD-MS^2^ in the Ion Trap with an isolation window of 1.6 *m/z*, calibrated charge-dependent ETD parameters, rapid scan mode, automatic scan and mass range, an ACG target of 50,000, and a maximum injection time of 100 ms; (3) EThcD-MS^2^ as performed for (2) except with 20% supplemental activation energy.

#### BAP1 knockout total protein expression samples

Liquid chromatography-mass spectrometry (LC-MS) analysis of native peptide fractions was carried out on a nanoflow LC system, EASY-nLC 1000, coupled to an Orbitrap Eclipse Tribrid mass spectrometer. 3 μL of each resuspended fraction was loaded onto a monolithic column (Capillary EX-Nano MonoCap C18 HighResolution 2000, 0.1 x 2000 mm, Merck, Darmstadt Germany) fitted with a silica coated PicoTip emitter (New Objective FS360-20-10-D). Solvents A and B were the same as described above for interactome samples. Peptides were separated over 180 min at a flow rate of 500 nL min^-1^ with the following gradient: 2-6% solvent B (10 min), 6-40% B (140 min), 40-98% B (1 min), and 98% B (29 min). MS^1^ spectra were acquired in the Orbitrap at 120K resolution with a scan range from 400-1500 *m/z*, an AGC target of 4 x 10^5^ and the maximum injection time set automatically. Features were filtered for monoisotopic peaks with a charge state of 2-5 and a minimum intensity of 5,000, with dynamic exclusion set to exclude features after 1 time for 60 s with a 10 ppm mass tolerance. Data-dependent MS^2^ analysis was performed using a top speed approach (cycle time of 5 s). Collision-induced dissociation (CID) fragmentation was performed with normalized collision energy of 35% and an activation time of 10 ms with an activation Q of 0.25 after quadrupole isolation of features using an isolation window of 0.7 *m/z*. Scans were acquired in the Ion Trap using automatic scan range, a maximum injection time of 250 ms, and rapid scan mode. HCD-SPS-MS^3^ scans were also performed in the Orbitrap on the top 10 SPS precursors with an isolation window of 0.7 *m/z*, 55% normalized collision energy, resolution of 60K, ACG target of 150,000, scan range of 100-500 *m/z*, and a maximum injection time of 118 ms. Precursor selection range, ion exclusion, and isobaric tag loss exclusion were all employed.

### MS data analysis

All raw files were searched using Proteome Discoverer v2.4.0.305 (Thermo Fisher Scientific) with the Byonic (Protein Metrics, Cupertino, CA) search node v3.7.4. Search parameter files are provided in **File S24**. Peak lists were searched against the species-specific complete UniProtKB databases and the UniprotKB/Swiss-Prot databases (*Mus musculus*, retrieved January 21, 2020; *Homo sapiens*, retrieved January 25, 2020) supplemented with a database of frequently observed contaminants (245 sequences). The Swiss-Prot only searches were used to match O-GlcNAc sites with known PTMs in the PhosphoSitePlus database and for the BAP1 KO quantitative O-GlcNAcomics and proteomics experiment to facilitate matching of O-GlcNAcylation peptides to their parent protein expression. The following parameters were set for all searches: MS^1^ tolerance of 5 ppm, MS^2^ tolerance of 10 ppm or 0.5 Da for Orbitrap and Ion Trap spectra, respectively, minimum peptide length of six amino acids, a maximum of three missed cleavages, carbamidomethylation of cysteine residues as a fixed modification, acetylation of the protein N terminus and oxidation of methionine (common 2) as variable modifications, and the tagged O-GlcNAc was as a variable modification (common 2), dimethylation or TMT labeling of lysines and peptide N-termini as fixed modifications where appropriate, and phosphorylation as a variable modification for the protein expression samples. Data was filtered to a 1% false discovery rate on PSMs using the Percolator or target decoy algorithm in Proteome Discoverer. Localization of O-GlcNAc sites was performed using the ptmRS node. For HCD spectra, equal localization probabilities were assigned to every serine/threonine due to neutral loss of the glycan. All raw files were uploaded to ProteomeXchange (PXD035902).

### Data analysis

#### General methods and statistics

All data analysis was performed using the Python language (Python Software Foundation, Python Language Reference, v3.7.3, available at http://www.python.org) with all Anaconda packages installed (Anaconda Software Distribution, v4.8.3, available at https://anaconda.com) unless otherwise noted. Information from the Uniprot Knowledgebase^91^ and PhosphoSitePlus^53^ database used for the integrated Cytoscape tables (**Data S1-S4**) was retrieved on April 22, 2020 and April 28, 2020, respectively. For the limma moderated t-test^133^ (package, limma v3.42.2) and inverted beta binomial tests^134^ (package, ibb v13.06), the R language (R Development Core Team, v3.6.1, available at https://www.R-project.org) was employed using the rpy2 (Laurent Gautier, v2.9.4) Python package. Original code is available in **File S27**. Multiple hypothesis testing correction was performed with either the Benjamini-Hochberg Method (inverted beta binomial testing) or Holm-Bonferroni method (western blotting). Fisher’s exact test was performed in GraphPad Prism (San Diego, CA, v9.3.0). In all cases, a *P* value of less than 0.05 was considered significant.

#### Bioinformatics analysis of OGT interactome and O-GlcNAcome

The input for the network analysis included (1) OGT interactors and substrates identified as above, and (2) known interactions from protein-protein interaction (PPI) databases. Two PPI databases were utilized: BioGRID (*Homo sapiens*-specific dataset, downloaded February 25, 2020) and IntAct (human-human PPIs, downloaded February 25, 2020 and filtered for only multivalidated interactions). Interactions between an OGT interactor and other interactors or substrates were filtered to generate the edges of the O-GlcNAc network. Edges from individual OGT interactor “baits” with more than thirty connections to other protein “prey” were removed to avoid biasing during community clustering. Examples of these proteins include ribosomal proteins, heterogeneous nuclear ribonucleoproteins (hnRNPs), and heat shock proteins (HSPs). Overall, these eliminated edges essentially coincided with interactions from proteins highly represented within the CRAPome,^135^ implying a high rate of non-specific interactions. However, edges generated in the reverse direction (e.g., from an interactor/substrate “bait” to a ribosomal protein “prey”) were retained. The nodes and edges were then used to build a network of OGT interactors and substrates, which was visualized and analyzed using Cytoscape v3.7.2.^57^ To partition the network, we used the Community cluster (GLay) algorithm^59^ in the Cytoscape plug-in clusterMaker2^60^ (v1.3.1), which is a community clustering algorithm that implements the Girvan Newman fast greedy algorithm.^58^

The protein names from the whole network or from each cluster were submitted for statistical overrepresentation testing using g:Profiler.^136^ Only terms with a *P* value of less than 0.01 with the g:SCS significance threshold were used, and electronic GO annotations were excluded. The complete datasets along with all submission parameters are provided in **Files S25** (HEK293T cells) and **S26** (brain and liver). To group GO terms and appropriately name clusters, the Cytoscape plugin ClueGO v2.5.6^65^ was used. For each cluster, the proteins were submitted to ClueGO analysis with GO Biological Processes (downloaded February 17, 2020) again with all electronic annotations excluded. GO Term Fusion was used, and only pathways with a *P* value of less than 0.01 were considered significant. Network specificity was set to medium with all dependent parameters left as default. GO Term grouping was employed.

### Neuronal culture, stimulation, and lysis

Primary mouse cortical neurons were harvested as previously described,^4^ plated on poly-D-lysine coated plates, and cultured in Neurobasal medium with 1% P/S, 2 mM GlutaMAX Supplement, and 1x B-27 Plus. Half of the media was changed every 2-4 days. After 20 days in vitro, neuronal activity was silenced using 10 μM tetrodotoxin (Tocris Biosciences, Bristol, United Kingdom) and 100 μM D-AP5 (Tocris Biosciences), and a subset was also treated with 50 μM Thiamet-G. The following day, silenced neurons were depolarized with 50 mM KCl or vehicle for 2 h and subsequently lysed with 2% SDS, 100 mM HEPES pH 7.9 containing cOmplete protease inhibitor cocktail and 100 μM Thiamet-G. Protein concentrations were measured using the BCA assay. For the immunoprecipitation experiments, silenced and depolarized neurons were prepared as described and lysed with 1% Triton X-100 in Tris-buffered saline (TBS) pH 7.6 containing cOmplete protease inhibitor cocktail, 100 μM Thiamet-G, and 0.25 U/μL benzonase nuclease (Santa Cruz Biotechnology).

### Chemoenzymatic labeling of neuronal lysates

O-GlcNAcylated proteins from cell lysates (150 μg) were labeled as previously described.^137^ Briefly, proteins were precipitated by the CH_3_OH/CHCl_3_/H_2_O method described above, and pellets were then resolubilized with 40 µL of 1% SDS and 20 mM HEPES pH 7.9 for 5 min at 95 °C. ddH_2_O (49 µL), 5.5 mM MnCl_2_ (11 μL), and 80 µL 2.5x GalT labeling buffer were added, and the solution was vortexed gently before adding 10 µL of 1 mg mL^-1^ Y289L GalT and 10 µL of 0.5 mM UDP-GalNAz in 10 mM HEPES pH 7.9. The reaction was then rotated end-over end at 4 °C overnight. Control experiments were carried out in parallel in the absence of UDP-GalNAz. The following day, samples were alkylated with 12.5 mM iodoacetamide for 1 h at room temperature protected from light. Proteins were then precipitated as before and resolubilized by boiling in 199.6 μL of 1% SDS and TBS pH 7.6. After allowing the redissolved proteins to cool to room temperature, 0.4 μL of 50 mM WS DBCO Biotin (Click Chemistry Tools) was added. The solution was rotated end-over-end for 1 h in the dark before being precipitated as described above. 10% of the reaction volume was taken as input. The remaining aliquot was then incubated with streptavidin magnetic beads (Thermo Fisher) for 1.5 h in the dark. Beads were then washed with 5 x 0.5 mL of low salt buffer (100 mM Na_2_HPO_4_, 150 mM NaCl, 0.1% SDS, 1% Triton X-100, and 0.5% sodium deoxycholate) and 5 x 1 mL of high salt buffer (100 mM Na_2_HPO_4_, 500 mM NaCl, and 0.2% Triton X-100). Biotinylated proteins were eluted by boiling the resin in 50 mM Tris-HCl pH 6.8, 2.5% SDS, 100 mM DTT, 10% glycerol, and 2 mM biotin for 15 min with intermittent vortexing.

### Western blotting

Primary antibodies were obtained from Cell Signaling Technology (Danvers, MA), and secondary antibodies were obtained from Thermo Fisher Scientific unless otherwise specified. Cell lysates were diluted with 4x SDS-PAGE buffer (200 mM Tris-HCl pH 6.8, 400 mM DTT, 8% SDS, 0.4% bromophenol blue, and 40% glycerol), resolved on a NuPAGE 4-12% Bis-Tris protein gel (Thermo Fisher Scientific), and transferred to Immobilion-FL PVDF membrane (MilliporeSigma). Blots were blocked for 1 h at room temperature with Odyssey Blocking Buffer (TBS) (LI-COR Biosciences, Lincoln NE) before probing overnight at 4 °C with the following antibodies diluted in blocking buffer at 1:1000 unless otherwise specified: anti-OGT (Proteintech, Rosemont, IL; 11576-2-AP), anti-HA (C29F4), anti-Nup62 (BD Biosciences, Franklin Lakes, NJ; 610498), anti-HCFC1 (Bethyl Laboratories, A301-399A-M), anti-SET1A (61702S), anti-RBBP5 (61702S), anti-BAP1 (Rabbit, Bethyl Laboratories, Montgomery, TX; A302-243A-M), anti-WDR5 (13105S), anti-FoxK1 (Bethyl Laboratories, A301-728A-M), anti-FoxK2 (12008S), anti-liprin-α1 (Proteintech, 14175-1-AP), anti-BAP1 (Mouse, Santa Cruz Biotechnology, sc-28383), or anti-α-tubulin (Sigma Aldrich, T9026, 1:3000). Blots were rinsed three times with TBS and 0.1% Tween 20 (TBST) and then probed with the appropriate secondary antibodies at 1:10,000 dilution in blocking buffer: anti-mouse AlexaFluor 680 conjugate (A21057), anti-rabbit AlexaFluor 680 (A21109), anti-rabbit AlexaFluor 790 (A11369), or anti-mouse IgG DyLight 800 (Thermo Fisher, A11357). Blots were washed 3 x 5 min with TBST and then imaged using an Odyssey Infrared Imaging System (LI-COR Biosciences). Images were processed using ImageStudio v5.2 (LI-COR Biosciences). To calculate O-GlcNAcylation stoichiometry, the ratios of eluents to inputs, weighted by proportion of the total protein in each lane, were calculated as previously described.^46^

### Co-immunoprecipitation in OGT-FH neurons

Neurons harvested from OGT-FH mice were grown, silenced, treated with KCl or vehicle, and lysed after 21 days in vitro as described above. From each condition, 1 mg of protein was diluted to 1 mL with TBS containing 100 μM Thiamet-G,and cOmplete protease inhibitor cocktail. 15 μg of protein was saved as input. The remaining lysate was incubated with anti-FLAG magnetic beads (MilliporeSigma) at 4 °C overnight with end-over-end rotation. The following day, beads were washed 3 times with 0.1% Triton X-100 in TBS (pH 7.9) and eluted by boiling in 2% SDS and TBS pH 8.0. Inputs and eluents were run on SDS-PAGE and western blotted for HA and liprin-α1 as described above.

## SUPPLEMENTAL INFORMATION TITLES AND LEGENDS

**Data S1. O-GlcNAc network of OGT interactors and substrates from HEK293T cells.**

**Data S2. O-GlcNAc network of OGT interactors and substrates from wild-type and BAP1 knockout HEK293T cells with O-GlcNAc site quantification.**

**Data S3. O-GlcNAc network of OGT interactors and substrates from forebrain tissue of OGT-FH mice.**

**Data S4. O-GlcNAc network of OGT interactors and substrates from liver tissue of OGT-FH mice.**

**File S1. OGT interactors identified from HEK293T cells.** limmaFC, limmapVal, limmaB: SILAC ratio log2 fold change, *P* value, and B-statistic, respectively, calculated using the limma moderated t-test. ibbFC, ibbpVal: spectral counts log2 fold change and associated *P* value after correction for multiple comparisons using the Benjamini-Hochberg method, respectively, calculated using the inverted beta binomial test.

**File S2. O-GlcNAc modification sites identified from HEK293T cells.** Region ID: unique identifier for each O-GlcNAc site or region. Min Sites: the minimum number of O-GlcNAc needed to explain the MS data (see also **Figure S2**). Site ID Constraints: amino acid residue(s) corresponding to each O-GlcNAc site; sites have only one possible residue, while regions have multiple possible residues (see also **Figure S2**).

**File S3. OGT substrates identified from HEK293T cells.** Entries contain Uniprot accession code, description, and gene names.

**File S4. Gene set enrichment analysis of OGT interactors, substrates, and individual network clusters from HEK293T cells.** *P* values reported for enrichment of each gene or protein group in the substrate dataset (“Substrates”), the interactor dataset (“Interactors”), the combined dataset (“All”), and partitioned clusters within the network (“Cluster1-10”). Tabs correspond to individual databases.

**File S5. Gene set newly enriched by network clustering**. *P* values reported for enrichment of each gene or protein group in the partitioned clusters within the network (“Cluster1-10”) that were not found in the substrate, interactor, or combined datasets. Tabs correspond to individual databases.

**File S6. OGT interactors and substrates from HEK293T cells that are involved in mRNA splicing.** Proteins from the HEK293T network were cross-referenced with entries in the Spliceosome Database. For each protein, its spliceosome complex(es), class or family annotations, and PDB structures are also provided.

**File S7. Post-translational modifications that are overlapping or proximal to identified O-GlcNAc sites in HEK293T cells.** Overlapping and proximal sites compiled from PhosphoSitePlus. Protein: Uniprot accession code; Site ID Constraint: amino acid site or region; Type: site or region, (Modification)Info: site of overlapping or neighboring modification as well as any associated information (e.g., “disease”, “regulatory”, “PTMVar - PTMs impacted by variants”). Tabs are separated by overlapping or within 10 amino acid residues and whether the Full Proteome or Swiss-Prot database were used.

**File S8. Quantification of protein domain, region, motif, and repeat annotations for OGT interactors, OGT substrates, and O-GlcNAc modification sites from HEK293T cells.** Annotations for each substrate, interactor, and individual site within HEK293T network were quantified and ranked.

**File S9. Network-based ranking of potential OGT adaptors from HEK293T cells.** Scores were calculated by the adaptor ranking algorithm based on HEK293T network connectivity.

**File S10. Quantified O-GlcNAc modification sites from wild-type and BAP1 knockout HEK293T cells.** See **File S2** for column annotations. ‘Experiment’ = which experimental run(s) in which the site or region was detected; logFC, P-Val, and B: log2 fold change, *P* value, and B statistic, respectively, calculated by TMT quantification; logFC protein: log2 fold change for protein containing each site or region; logFC_Corrected, P-Val_Corrected, B_Corrected: log2 fold change, *P* value, and B-statistic, respectively, for each site corrected for individual protein quantification.

**File S11. OGT interactors identified from forebrain tissue of OGT-FH mice.** See **File S1** for annotations of columns.

**File S12. OGT interactors identified from liver tissue of OGT-FH mice.** See **File S1** for annotations of columns.

**File S13. O-GlcNAc modification sites identified from forebrain tissue of OGT-FH mice.** See **File S2** for column annotations.

**File S14. O-GlcNAc modification sites identified from liver tissue of OGT-FH mice.** See **File S2** for column annotations.

**File S15. OGT substrates identified from forebrain tissue of OGT-FH mice.** Entries contain Uniprot accession code, description, and gene names.

**File S16. OGT substrates identified from liver tissue of OGT-FH mice.** Entries contain Uniprot accession code, description, and gene names.

**File S17. Gene set enrichment analysis of OGT interactors, substrates, and individual network clusters from forebrain and liver tissue of OGT-FH mice. See** File S4 for column annotations.

**File S18. Network-based ranking of potential OGT adaptors from forebrain tissue of OGT FH mice.** Scores were calculated by the adaptor ranking algorithm based on brain network connectivity.

**File S19. Network-based ranking of potential OGT adaptors from liver tissue of OGT-FH mice.** Scores were calculated by the adaptor ranking algorithm based on liver network connectivity.

**File S20. Post-translational modifications that are overlapping or proximal to identified O-GlcNAc sites in forebrain tissue of OGT-FH mice. See** File S7 for column annotations.

**File S21. Post-translational modifications that are overlapping or proximal to identified O-GlcNAc sites in liver tissue of OGT-FH mice.** See **File S7** for column annotations.

**File S22. OGT interactors and substrates from forebrain tissue of OGT-FH mice that are involved in synaptic function.** Proteins in the brain network were queried for “SYNAP*” in their associated functional annotation data and compiled.

**File S23. Quantification of protein domain, region, motif, and repeat annotations for OGT interactors, OGT substrates, and O-GlcNAc modification sites from forebrain and liver tissue of OGT-FH mice.** See File **S8**.

**File S24. Search parameter files for MS data using Proteome Discoverer with Byonic search node.**

**File S25. Complete dataset and submission parameters for g:Profiler analysis of proteins in the HEK293T network.**

**File S26. Complete dataset and submission parameters for g:Profiler analysis of proteins in the murine forebrain and liver networks.**

**File S27. Original Python code to perform sites and region analysis, network generation, and adaptor protein ranking.**

