## Supplemental Information for "Functional glycoproteomics by integrated network assembly and partitioning"

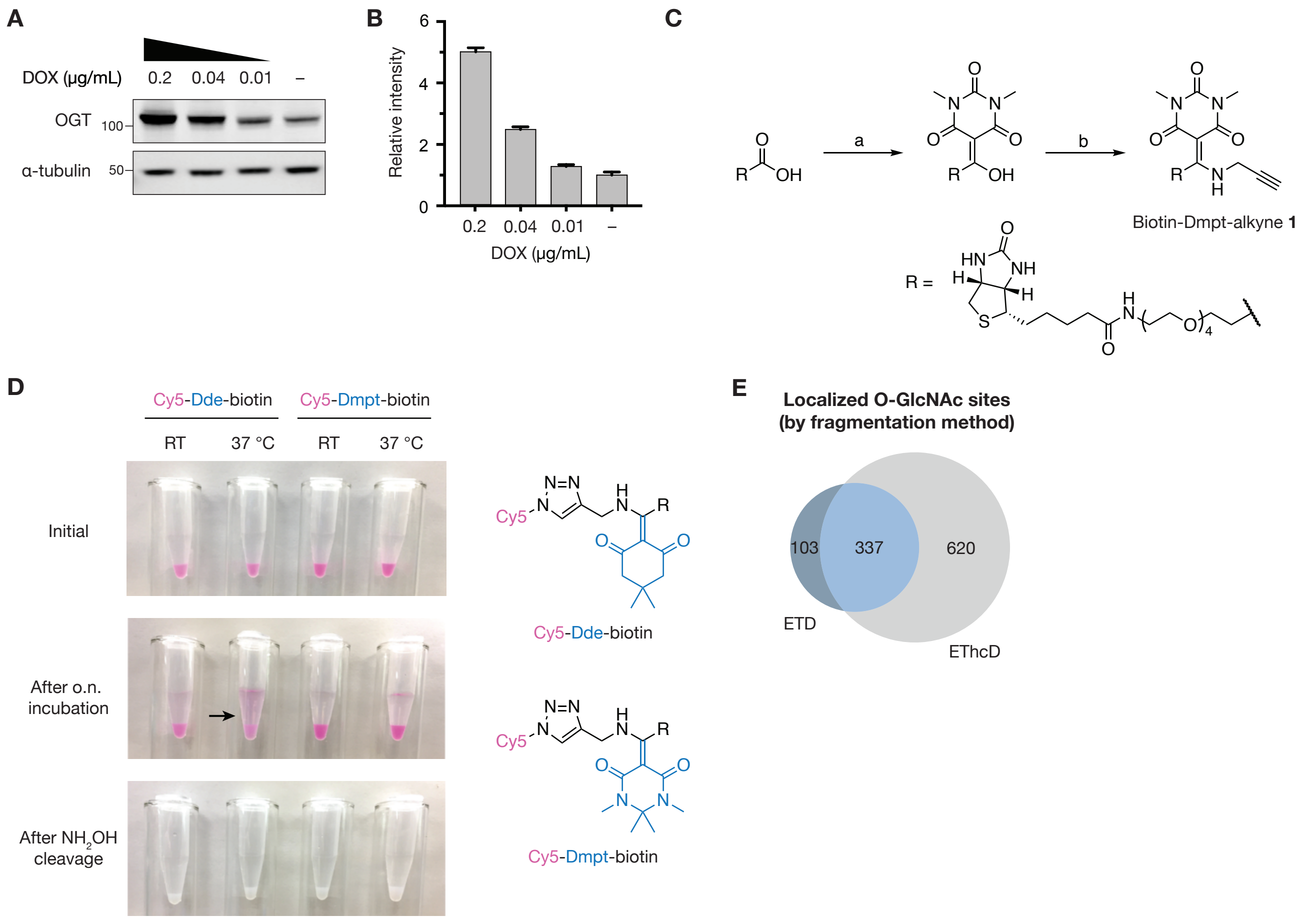

A
DOX (µg/mL)
0.2
0.04
0.01
–
OGT
100
α-tubulin
50
B
Relative intensity
DOX (µg/mL)
C
a
b
Biotin-Dmpt-alkyne 1
R =
D
Cy5-Dde-biotin
Cy5-Dmpt-biotin
RT
37 °C
Initial
After o.n. incubation
After NH2OH cleavage
E
Localized O-GlcNAc sites (by fragmentation method)
103
337
620
ETD
EThcD

**Figure S1. Novel methods for expression and characterization of OGT, related to Figure 2.**

(A-B) Western blotting and quantification of inducible OGT-FH expression by HEK293T cells. Cells were treated with the indicated doxycycline (DOX) concentrations for 24 h. After lysis, lysates were probed for OGT-FH expression by anti-HA western blotting. Quantification was performed by normalization to α-tubulin expression.

(C) Synthesis of biotin-Dmpt-alkyne **1**. (a) DBA (1.3 eq), EDC·HCl (1.3 eq), TEA (1.5 eq), DMAP (0.1 eq), DCM, 0 °C to RT, 16 h, 92% yield. (b) Propargylamine (neat), 50 °C, 4 h, 28% yield. Detailed synthetic protocols are provided in the Methods section.

(D) Comparative stability of biotin-Dde-alkyne and biotin-Dmpt-alkyne. (i) Biotin-Dde-alkyne or biotin-Dmpt-alkyne **1** was reacted with azido-Cy3 and immobilized onto high-capacity Neutravidin resin. After overnight incubation in phosphate-buffered saline, the dye conjugated to the Dde linker showed leaching into the buffer at higher temperatures necessary for tryptic digestion (arrow). Both linkers showed efficient release of the immobilized dye after treatment with $NH_2OH$.

(E) Pie chart of the overlap between all unique, localized O-GlcNAc sites identified by ETD and EThcD fragmentation methods in HEK293T cells.

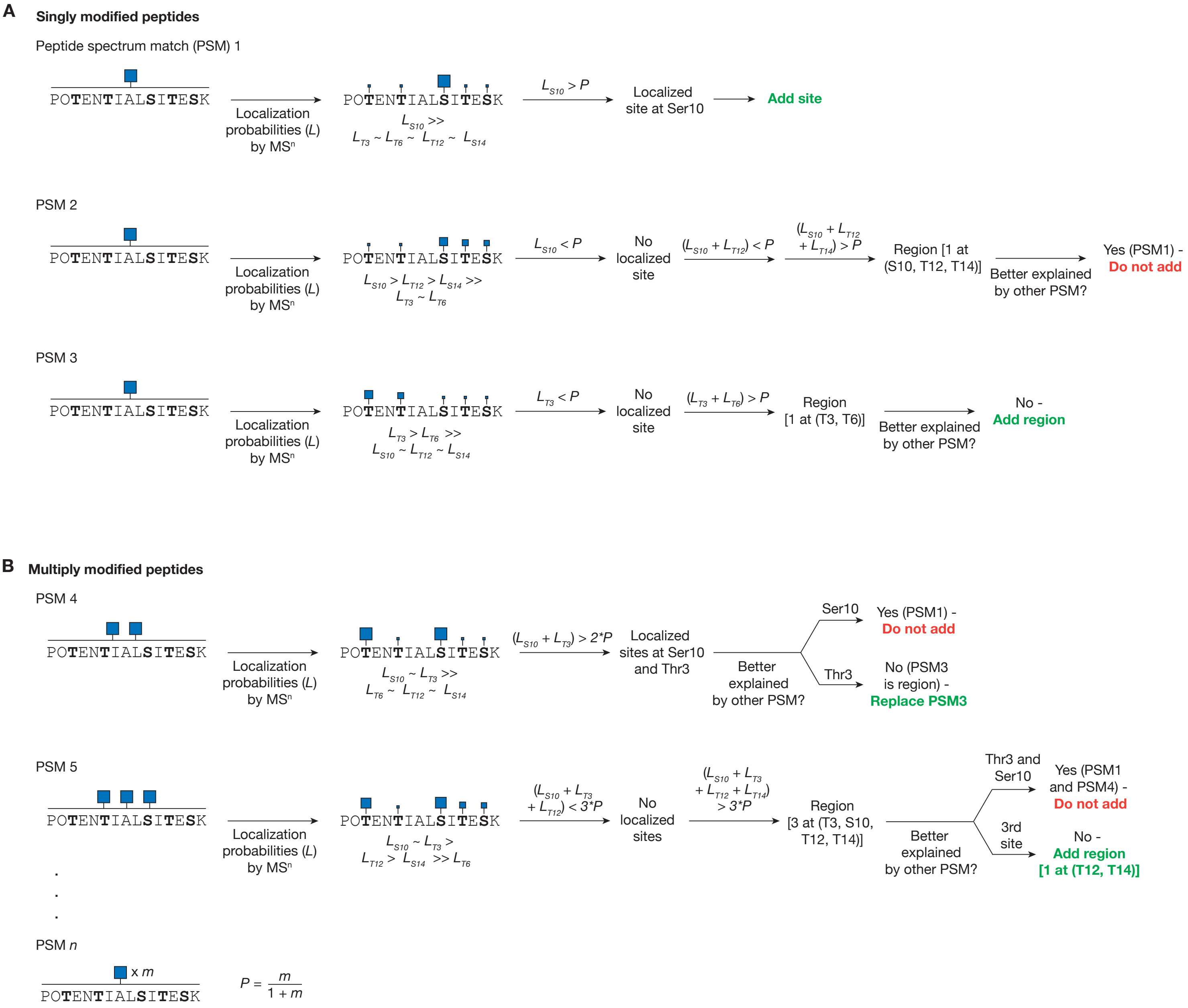
A
Singly modified peptides
Peptide spectrum match (PSM) 1
POTENTIALSITESK
Localization probabilities (L) by MSn
LS10 >> LT3 ~ LT6 ~ LT12 ~ LS14
LS10 > P
Localized site at Ser10
Add site
PSM 2
POTENTIALSITESK
Localization probabilities (L) by MSn
LS10 > LT12 > LS14 >> LT3 ~ LT6
LS10 < P
No localized site
(LS10 + LT12) < P
(LS10 + LT12 + LT14) > P
Region [1 at (S10, T12, T14)]
Better explained by other PSM?
Yes (PSM1) -
Do not add
PSM 3
POTENTIALSITESK
Localization probabilities (L) by MSn
LT3 > LT6 >> LS10 ~ LT12 ~ LS14
LT3 < P
No localized site
(LT3 + LT6) > P
Region [1 at (T3, T6)]
Better explained by other PSM?
No -
Add region
B
Multiply modified peptides
PSM 4
POTENTIALSITESK
Localization probabilities (L) by MSn
LS10 ~ LT3 >> LT6 ~ LT12 ~ LS14
(LS10 + LT3) > 2*P
Localized sites at Ser10 and Thr3
Better explained by other PSM?
Ser10
Yes (PSM1) -
Do not add
Thr3
No (PSM3 is region) -
Replace PSM3
PSM 5
POTENTIALSITESK
Localization probabilities (L) by MSn
LS10 ~ LT3 > LT12 > LS14 >> LT6
(LS10 + LT3 + LT12) < 3*P
No localized sites
(LS10 + LT3 + LT12 + LT14) > 3*P
Region [3 at (T3, S10, T12, T14)]
Better explained by other PSM?
Thr3 and Ser10
Yes (PSM1 and PSM4) -
Do not add
3rd site
No -
Add region
[1 at (T12, T14)]
PSM n
x m
POTENTIALSITESK
P = m / (1 + m)

**Figure S2. Workflow to identify O-GlcNAc sites and regions, related to Figure 2.**

Method to process peptide spectrum matches (PSMs) and compile a total list of the O-GlcNAc localized sites and unlocalized regions within a dataset. O-GlcNAc sites are only considered localized if the localization probability (ptmRS or *L*) is greater than random chance. The threshold for nonrandom probability that a site is modified (*P*) is calculated as the number of sites in the PSM (*m*) divided by 1 + *m*.

(A) For a singly modified PSM, localization probabilities are calculated for each site based on $MS^n$ fragmentation and ranked in descending order. The highest ranked probability is compared to *P*. If $L > P$ as in PSM1, then the site is considered localized and added to the total list as a localized site. If $L < P$ as in PSM2 and PSM3, then probabilities are summed in descending order until the sum is greater than *P*. This new region is then compared to other sites within the list to see if it is better explained by another PSM. For PSM2, the region is better explained by a localized site in PSM1, so it is not added to the list. For PSM3, the region is not better explained by another PSM, so it is added to the list as a region.

(B) For a PSM modified by *m* sites, the highest *m* localization probabilities are summed and compared against $m*P$. For PSM4, this approach yields two potential O-GlcNAc sites, which are then compared with the existing list. One site is already included on the list from PSM1 and is not included to the list as a new entry. The other site better explains the region from PSM3 and is included to the list as a replacement for the PSM3 region. For PSM5, the workflow generates a region of three sites, two of which are better explained by other PSMs. Therefore, PSM5 only adds a single region to the overall list. For PSMs where multiple amino acid residues have equal localization probability such as HCD spectra that lack localization, a region incorporating all sites is added to the total list unless the site is better explained by another PSM.

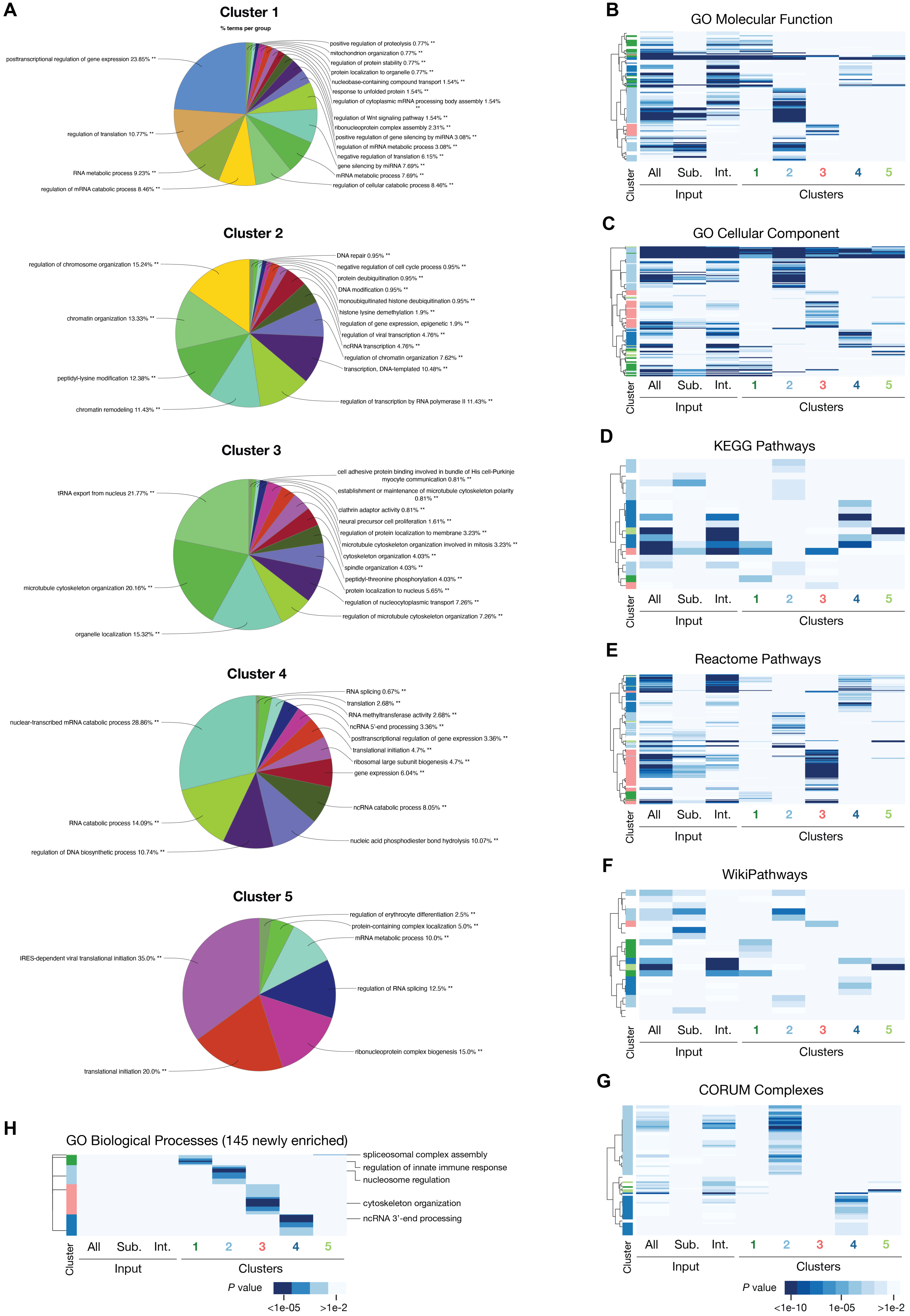

A
Cluster 1
% terms per group
posttranscriptional regulation of gene expression 23.85% **
regulation of translation 10.77% **
RNA metabolic process 9.23% **
regulation of mRNA catabolic process 8.46% **
positive regulation of proteolysis 0.77% **
mitochondrion organization 0.77% **
regulation of protein stability 0.77% **
protein localization to organelle 0.77% **
nucleobase-containing compound transport 1.54% **
response to unfolded protein 1.54% **
regulation of cytoplasmic mRNA processing body assembly 1.54% **
regulation of Wnt signaling pathway 1.54% **
ribonucleoprotein complex assembly 2.31% **
positive regulation of gene silencing by miRNA 3.08% **
regulation of mRNA metabolic process 3.08% **
negative regulation of translation 6.15% **
gene silencing by miRNA 7.69% **
mRNA metabolic process 7.69% **
regulation of cellular catabolic process 8.46% **
Cluster 2
regulation of chromosome organization 15.24% **
chromatin organization 13.33% **
peptidyl-lysine modification 12.38% **
chromatin remodeling 11.43% **
DNA repair 0.95% **
negative regulation of cell cycle process 0.95% **
protein deubiquitination 0.95% **
DNA modification 0.95% **
monoubiquitinated histone deubiquitination 0.95% **
histone lysine demethylation 1.9% **
regulation of gene expression, epigenetic 1.9% **
regulation of viral transcription 4.76% **
ncRNA transcription 4.76% **
regulation of chromatin organization 7.62% **
transcription, DNA-templated 10.48% **
regulation of transcription by RNA polymerase II 11.43% **
Cluster 3
tRNA export from nucleus 21.77% **
microtubule cytoskeleton organization 20.16% **
organelle localization 15.32% **
cell adhesive protein binding involved in bundle of His cell-Purkinje myocyte communication 0.81% **
establishment or maintenance of microtubule cytoskeleton polarity 0.81% **
clathrin adaptor activity 0.81% **
neural precursor cell proliferation 1.61% **
regulation of protein localization to membrane 3.23% **
microtubule cytoskeleton organization involved in mitosis 3.23% **
cytoskeleton organization 4.03% **
spindle organization 4.03% **
peptidyl-threonine phosphorylation 4.03% **
protein localization to nucleus 5.65% **
regulation of nucleocytoplasmic transport 7.26% **
regulation of microtubule cytoskeleton organization 7.26% **
Cluster 4
nuclear-transcribed mRNA catabolic process 28.86% **
RNA catabolic process 14.09% **
regulation of DNA biosynthetic process 10.74% **
RNA splicing 0.67% **
translation 2.68% **
RNA methyltransferase activity 2.68% **
ncRNA 5'-end processing 3.36% **
posttranscriptional regulation of gene expression 3.36% **
translational initiation 4.7% **
ribosomal large subunit biogenesis 4.7% **
gene expression 6.04% **
ncRNA catabolic process 8.05% **
nucleic acid phosphodiester bond hydrolysis 10.07% **
Cluster 5
IRES-dependent viral translational initiation 35.0% **
translational initiation 20.0% **
regulation of erythrocyte differentiation 2.5% **
protein-containing complex localization 5.0% **
mRNA metabolic process 10.0% **
regulation of RNA splicing 12.5% **
ribonucleoprotein complex biogenesis 15.0% **
B
GO Molecular Function
C
GO Cellular Component
D
KEGG Pathways
E
Reactome Pathways
F
WikiPathways
G
CORUM Complexes
Cluster
All
Sub.
Int.
Input
1
2
3
4
5
Clusters
H
GO Biological Processes (145 newly enriched)
spliceosomal complex assembly
regulation of innate immune response
nucleosome regulation
cytoskeleton organization
ncRNA 3'-end processing
P value
<1e-05
>1e-2
<1e-10
1e-05

**Figure S3. Analysis of O-GlcNAc functional networks in HEK293T cells, related to Figure 3.**

(A) Pie charts of significantly enriched GO biological process terms for each annotated community. Pie charts were generated by ClueGO.

(B-G) Hierarchical clustering of functional annotations from OGT interactors (int.), substrates (sub.), and network clusters.

(H) Hierarchical clustering of 145 GO Biological Process terms that were newly enriched after network clustering. GO term with the lowest *P* value indicated on the right. ncRNA = noncoding RNA.

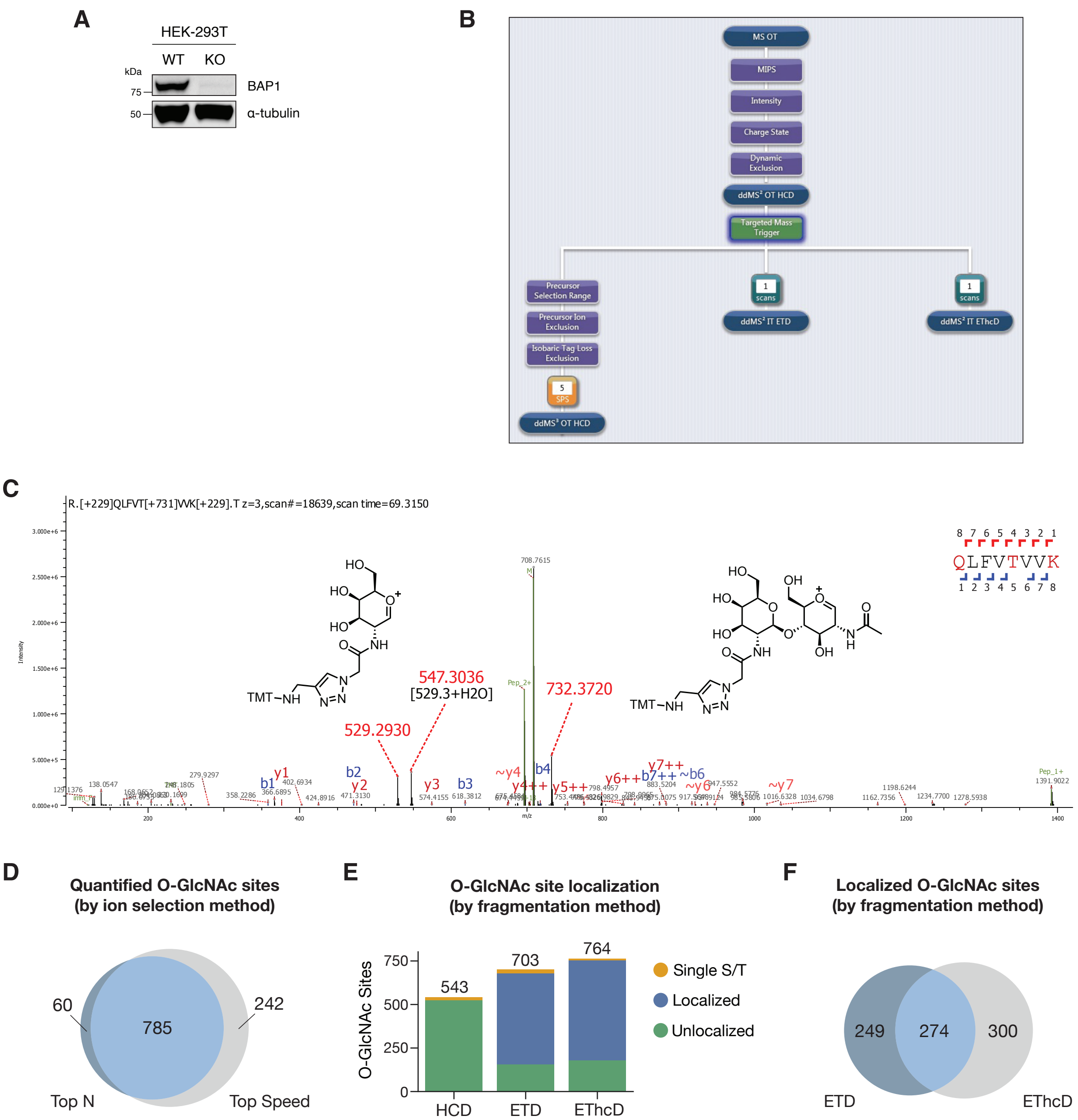

A
HEK-293T
WT KO
kDa
75
50
BAP1
α-tubulin
B
MS OT
MIPS
Intensity
Charge State
Dynamic Exclusion
ddMS² OT HCD
Targeted Mass Trigger
Precursor Selection Range
Precursor Ion Exclusion
Isobaric Tag Loss Exclusion
5 SPS
ddMS³ OT HCD
1 scans
ddMS² IT ETD
1 scans
ddMS² IT EThcD
C
R.[+229]QLFVT[+731]VVK[+229].T z=3,scan#=18639,scan time=69.3150
Intensity
m/z
8 7 6 5 4 3 2 1
QLFVTVVK
1 2 3 4 5 6 7 8
529.2930
547.3036
[529.3+H2O]
732.3720
708.7615
TMT−NH
1391.9022
D
Quantified O-GlcNAc sites
(by ion selection method)
60
785
242
Top N
Top Speed
E
O-GlcNAc site localization
(by fragmentation method)
O-GlcNAc Sites
750
500
250
0
543
703
764
HCD
ETD
EThcD
Single S/T
Localized
Unlocalized
F
Localized O-GlcNAc sites
(by fragmentation method)
249
274
300
ETD
EThcD

**Figure S4. Workflow and characterization of O-GlcNAc site quantification in wild-type and BAP1 knockout HEK293T cells, related to Figure 4.**

(A) Validation of BAP1 knockout in HEK293T cells. After lysis, lysates were probed for BAP1 expression by anti-BAP1 western blotting. Lysate loading was verified by anti-α-tubulin western blotting.

(B) Instrument workflow for O-GlcNAc site identification and quantification. After data-dependent HCD fragmentation with Orbitrap detection (ddMS$^2$ OT HCD), scans containing diagnostic ions were subjected to ddMS$^3$ OT HCD fragmentation for isobaric tag quantification or ETD/EThcD fragmentation with ion trap (ddMS$^2$ IT ETD/EThcD) for site localization.

(C) Representative MS scan of diagnostic ions and chemical structures produced by TMT-labeled chemoenzymatic linker fragment at 529.2930, 547.3036, and 732.3720 *m/z*.

(D) Analysis of localized and quantified O-GlcNAc sites. Pie chart shows the number of quantified O-GlcNAc sites based on ion selection method.

(E) Bar graph of localized sites produced by the three fragmentation methods used.

(F) Pie chart showing the overlap between unique, localized O-GlcNAc sites identified by ETD and EThcD fragmentation methods in the BAP1 KO quantitative glycomics experiment.

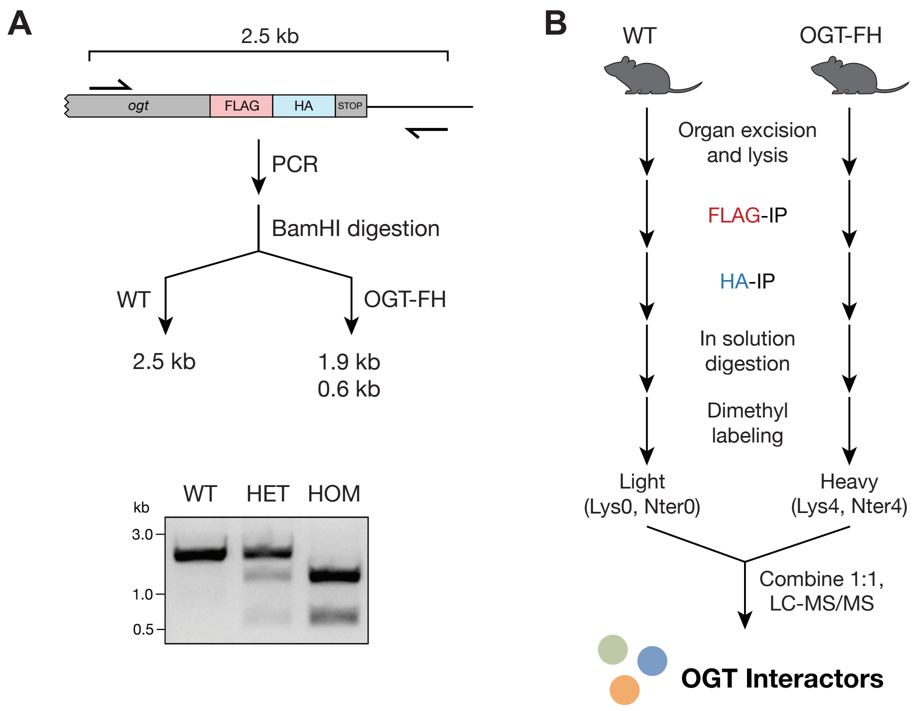

A
2.5 kb
ogt
FLAG
HA
STOP
PCR
BamHI digestion
WT
OGT-FH
2.5 kb
1.9 kb
0.6 kb
kb
WT
HET
HOM
3.0
1.0
0.5
B
WT
OGT-FH
Organ excision and lysis
FLAG-IP
HA-IP
In solution digestion
Dimethyl labeling
Light (Lys0, Nter0)
Heavy (Lys4, Nter4)
Combine 1:1, LC-MS/MS
OGT Interactors

**Figure S5. Workflow to characterize OGT-FH mouse model and in vivo interactors, related to Figure 5.**

(A) PCR scheme to genotype OGT-FH mouse model with representative agarose gel. A region containing the C-terminus of the *ogt* gene was amplified by PCR, and the PCR product was treated with BamHI. Wild-type animals yield a single band at 2.5 kb, and OGT-FH animals yield two bands at 1.9 kb and 0.6 kb.

(B) Workflow to identify in vivo OGT interactors. Organs were harvested and lysed in nondenaturing lysis buffer. Lysates were subjected to tandem FLAG and HA co-immunoprecipitation, and eluted proteins were subjected to in solution digestion. Tryptic peptides were tagged by dimethyl labeling using $CH_2O$ or $CD_2O$ and $NaBH_3CN$. For samples labeled with $CD_2O$, peptides gained 4 or 8 Da based on whether the terminal residue is Arg or Lys, respectively. Samples were then combined 1:1 and subjected to LC-MS/MS identification and quantification.

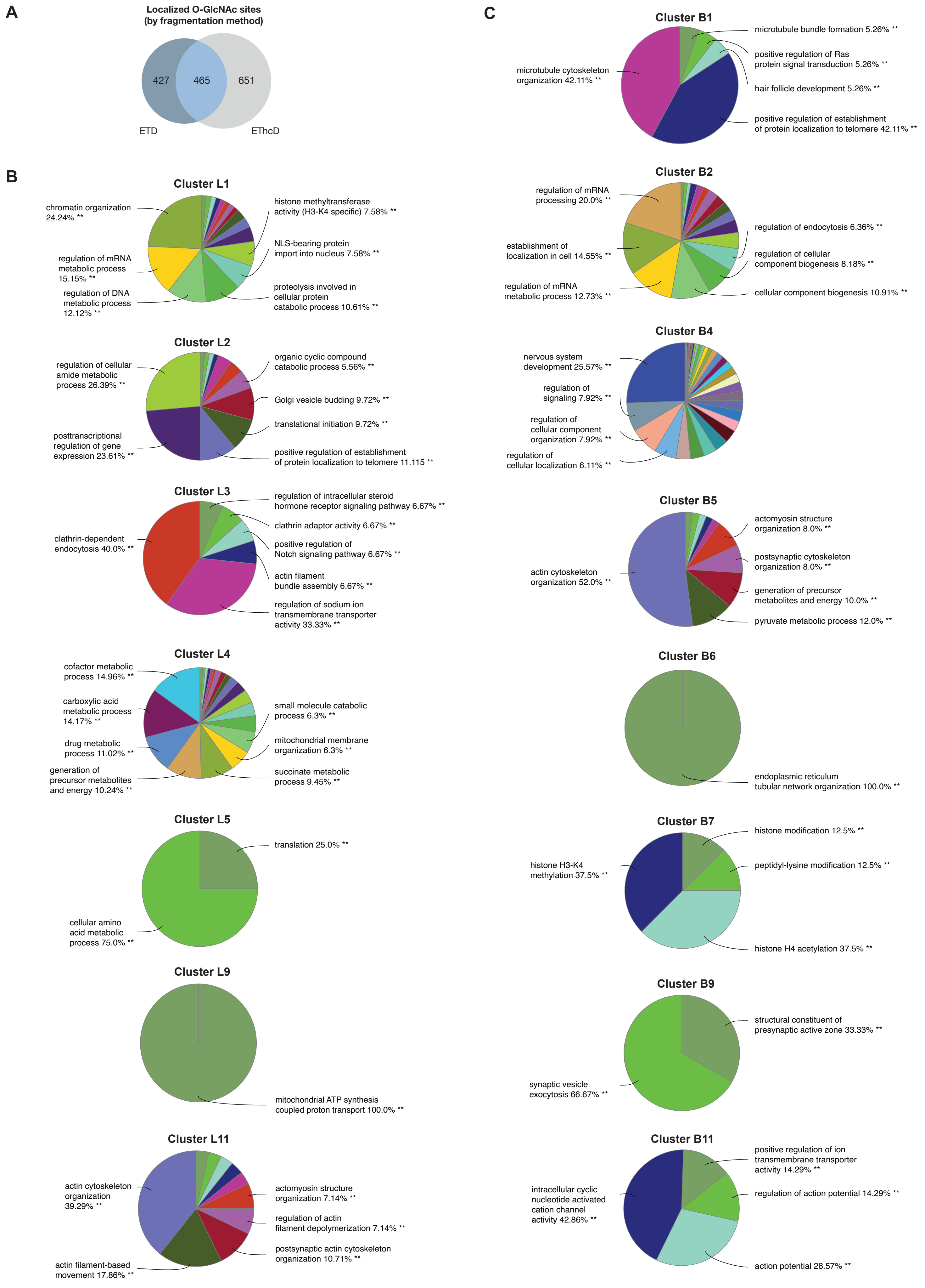
A
Localized O-GlcNAc sites
(by fragmentation method)
427
465
651
ETD
EThcD
B
Cluster L1
chromatin organization 24.24% **
regulation of mRNA metabolic process 15.15% **
regulation of DNA metabolic process 12.12% **
histone methyltransferase activity (H3-K4 specific) 7.58% **
NLS-bearing protein import into nucleus 7.58% **
proteolysis involved in cellular protein catabolic process 10.61% **
Cluster L2
regulation of cellular amide metabolic process 26.39% **
posttranscriptional regulation of gene expression 23.61% **
organic cyclic compound catabolic process 5.56% **
Golgi vesicle budding 9.72% **
translational initiation 9.72% **
positive regulation of establishment of protein localization to telomere 11.115 **
Cluster L3
clathrin-dependent endocytosis 40.0% **
regulation of intracellular steroid hormone receptor signaling pathway 6.67% **
clathrin adaptor activity 6.67% **
positive regulation of Notch signaling pathway 6.67% **
actin filament bundle assembly 6.67% **
regulation of sodium ion transmembrane transporter activity 33.33% **
Cluster L4
cofactor metabolic process 14.96% **
carboxylic acid metabolic process 14.17% **
drug metabolic process 11.02% **
generation of precursor metabolites and energy 10.24% **
small molecule catabolic process 6.3% **
mitochondrial membrane organization 6.3% **
succinate metabolic process 9.45% **
Cluster L5
translation 25.0% **
cellular amino acid metabolic process 75.0% **
Cluster L9
mitochondrial ATP synthesis coupled proton transport 100.0% **
Cluster L11
actin cytoskeleton organization 39.29% **
actin filament-based movement 17.86% **
actomyosin structure organization 7.14% **
regulation of actin filament depolymerization 7.14% **
postsynaptic actin cytoskeleton organization 10.71% **
C
Cluster B1
microtubule cytoskeleton organization 42.11% **
microtubule bundle formation 5.26% **
positive regulation of Ras protein signal transduction 5.26% **
hair follicle development 5.26% **
positive regulation of establishment of protein localization to telomere 42.11% **
Cluster B2
regulation of mRNA processing 20.0% **
establishment of localization in cell 14.55% **
regulation of mRNA metabolic process 12.73% **
regulation of endocytosis 6.36% **
regulation of cellular component biogenesis 8.18% **
cellular component biogenesis 10.91% **
Cluster B4
nervous system development 25.57% **
regulation of signaling 7.92% **
regulation of cellular component organization 7.92% **
regulation of cellular localization 6.11% **
Cluster B5
actin cytoskeleton organization 52.0% **
actomyosin structure organization 8.0% **
postsynaptic cytoskeleton organization 8.0% **
generation of precursor metabolites and energy 10.0% **
pyruvate metabolic process 12.0% **
Cluster B6
endoplasmic reticulum tubular network organization 100.0% **
Cluster B7
histone H3-K4 methylation 37.5% **
histone modification 12.5% **
peptidyl-lysine modification 12.5% **
histone H4 acetylation 37.5% **
Cluster B9
structural constituent of presynaptic active zone 33.33% **
synaptic vesicle exocytosis 66.67% **
Cluster B11
intracellular cyclic nucleotide activated cation channel activity 42.86% **
positive regulation of ion transmembrane transporter activity 14.29% **
regulation of action potential 14.29% **
action potential 28.57% **

**Figure S6. Analysis of O-GlcNAc functional networks in murine forebrain and liver, related to Figure 6.**

(A) Pie chart showing the overlap between unique, localized O-GlcNAc sites identified by ETD and EThcD fragmentation methods in the combined liver and brain datasets.

(B-C) Pie charts of significantly enriched GO biological process terms for each annotated community in the liver and brain networks, respectively. Pie charts were generated by ClueGO.
